# RNA binding limits the ligand induced transcriptional potential of estrogen receptor-alpha (ERα)

**DOI:** 10.1101/2023.08.10.552751

**Authors:** Deepanshu Soota, Bharath Saravanan, Rajat Mann, Tripti Kharbanda, Dimple Notani

## Abstract

Transcription factors (TFs) primarily regulate gene expression by binding to DNA through their DNA binding domains (DBD). Additionally, approximately half of these TFs also interact with RNA. However, the role of RNA in enabling TF binding on chromatin and subsequent transcription is poorly understood. Estrogen receptor-α (ERα) is one such TF that activates genes in response to estrogen stimulation. Here, we report that ERα interacts with various types of RNAs in ligand dependent manner via its RNA binding motif in hinge region. RNA binding defects lead to a global loss of ERα binding in the genome, particularly at weaker ERα motifs. In the absence of RNA binding, the ERα exhibits dynamic behavior in the nucleus and unexpectedly, the dynamic binding coincides with robust polymerase loading on ERα bound chromatin regions. The higher occupancy of PolII was recapitulated by robust ligand induced transcription of ERα-regulated genes. Collectively, our results suggest that RNA interactions strengthen ERα binding to chromatin limiting the ligand-dependent transcriptional upregulation of estrogen-induced genes.

## Introduction

Transcription factors (TFs) are known for their ability to recognize and bind specific DNA motifs through their DNA binding domains (DBD). In addition to DNA binding, TFs also possess RNA binding capability therefore; they are DNA and RNA binding proteins (Cassiday and Maher III 2002; Hudson and Ortlund 2014). The role of this TF-RNA interaction has been extensively studied at various post-transcriptional levels, such as RNA splicing (Auboeuf et al. 2002; Carnesecchi et al. 2021; Girardot et al. 2018; Xu et al. 2021), RNA stability (Dhawan et al. 2007) and translational regulation (Poon and Chen 2008; Rödel et al. 2013; Tournillon et al. 2017; Xu et al. 2021). TFs interact with RNA via their DBD, canonical RNA binding motifs (RGG), Arginine Rich Motifs (ARM) or low complexity regions (Mann and Notani 2023). The RNA binding through DBD sequesters TF from binding to its cognate DNA motif (Ahmed et al. 2021; Dickey and Pyle 2017; Hamilton et al. 2022; Holmes et al. 2020; Hudson and Ortlund 2014; Pelham and Brown 1980; Yang et al. 2020).Conversely, RNA interactions via non-DBD regions can modulate the TF’s binding dynamics on chromatin that leads to transcriptional dysregulation (Clemens et al. 1993; Hou et al. 2020; Oksuz et al. 2023; Sigova et al. 2015; Xu and Koenig 2004; Yang et al. 2013; Yoshida et al. 2004).

Estrogen receptor-alpha (ERα) is a ligand dependent steroid hormone receptor which regulates estrogen induced genes by recruiting polymerase complexes on specific enhancers and promoters (Li et al. 2013). ERα interacts with specific RNAs that influence its recruitment on genome (Aiello et al. 2016; Lanz et al. 1999; Xue et al. 2016). Similar to other TFs, ERα also interacts with these RNA via its DBD (Yang et al. 2020) or via its hinge region that is connected to the DBD. In our study we have primarily focused on the hinge region of ERα, which harbors Arginine rich motif (ARM), a well-known RNA binding motif. Mutations in the ERα-ARM have been shown to drastically decrease RNA binding (Oksuz et al. 2023; Steiner et al. 2022; Xu et al. 2021). The point mutation in ARM (R269C) caused the loss of RNA binding ability and exhibited the down regulation of Tat-mediated trans-activation of reporter gene (Oksuz et al. 2023). This suggests that RNA binding increases trans-activation potential of TF for the better transcription. Conversely, the mutation in RRGG exhibited no effect on ERα binding to chromatin and transcription under basal signaling (Xu et al. 2021). Non-reliance of ERα on RNA could be due to lack of ligand stimulation in this study as the ligand modulates the binding of ERα on DNA and facilitates the recruitment of robust transcription machinery for target gene activation (Shang et al. 2000) suggesting, ERα:RNA interaction and its function may be influenced by the ligand-dependent events.

Therefore, we utilized estrogen stimulation to examine the nuclear role of ERα:RNA interaction. We performed fRIP-seq, biochemical fractionations and genome-wide studies on WT and RBM mutant (RRGG mutant) of ERα in MCF-7 cells upon estrogen stimulation. Through fRIP-seq analysis, we identified ERα interactions with RNA transcribing from diverse genomic regions. Further, we observed that the RNA binding-deficient mutant of ERα exhibits defects in DNA binding potential upon ligand stimulation, contrary to previous reports under basal signaling (Xu et al. 2021). Thus, the loss of RNA binding ability results in diminished genomic occupancy of ERα under signaling dependent context. Additionally, RNase A treatment followed by extraction of chromatin with and without the presence of nucleoplasmic fraction confirms the requirement of RNA for the stabilization of ERα on chromatin. Further, FRAP and biochemical fractionations revealed a dynamic binding of ERα to chromatin. Unexpectedly, we observed that this dynamic binding of ERα leads to increased RNA polymerase II loading and subsequent transcription of E2-regulated genes. Overall, our results suggest that RNA binding allows stable recruitment of TF on chromatin that limits the transcriptional potential of TF.

## Results

### ERα interacts with RNA

ERα interacts with mRNA under basal signaling (Joyce et al. 2010; Xu et al. 2021) and to specific lncRNAs (Aiello et al. 2016; Horie et al. 2022; Xue et al. 2016). To identify the RNA that interact with ERα after ligand stimulation, we performed fRIP-seq (Hendrickson et al. 2016) after 1 hour of estradiol (E2) treatment in MCF-7 cells. The enriched RNAs were from various types of regions in the genome (Fig. 1A). The highest proportions of RNA that interacted with ERα were introns (approx. 30%) and promoter or promoter-proximal regions (approx. 22%) followed by 3’UTR, exon and intergenic RNA. Fold changes over input exhibited 3’UTR followed by intergenic and intronic region to be highly enriched (Fig. 1B, C and Fig. S1A). Similar to genic regions, ERα exhibited interaction with eRNAs as shown for TFF1e (Fig. 1D). Since the fRIP-seq does not distinguish the RNA from chromatin or nucleoplasmic fraction, we intersected the ERα fRIP-seq peaks with RNA-seq from chromatin or nucleoplasmic fractions in MCF-7 cells (Ntini et al. 2018). ERα fRIP-seq peaks exhibited no preferential biases towards the RNA from specific nuclear fraction (Fig. 1E). As expected, the introns and distal intergenic regions were more enriched in chromatin fraction whereas; exons were enriched in nucleoplasmic fraction (Fig. S1B).

**Figure 1.**
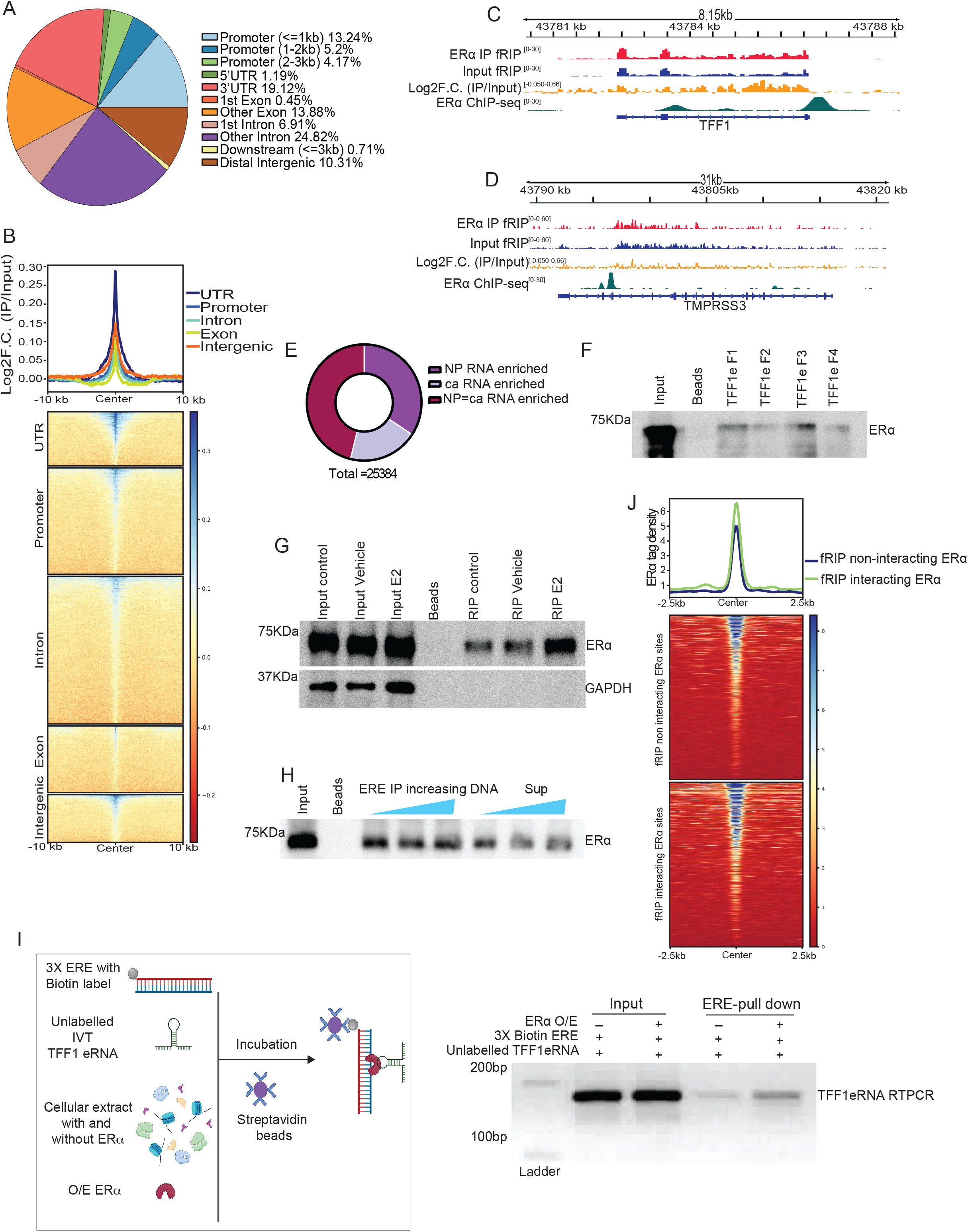
ERα interacts with RNA. A. Pie chart showing the genomic distribution of ERα interacting RNA by formaldehyde assisted RNA immunoprecipitation sequencing (fRIP-seq). B. Heatmap showing the enrichment of fRIP-seq immunoprecipitation over the input across categories of ERα interacting RNA. C. Genome browser screenshot showing fRIP-seq IP, Input, Log2F.C.(IP/Input) and ERα ChIP-seq for TFF1mRNA. D. Genome browser screenshot with fRIP-seq IP, Input, Log2F.C.(IP/Input) and ERα ChIP-seq for TFF1 eRNA. E. Proportions of ERα interacting RNA inside the nucleus (NP: Nucleoplasmic enriched RNA and ca: Chromatin associated RNA). F. Immunoblot for ERα on RNA pulldowns using biotin labelled TFF1 eRNA fragments. G. Immunoblot for ERα and GAPDH on RNA pulldowns using biotin labelled TFF1 eRNA with lysates from cells grown in DMEM or stripping media treated with either Vehicle or E2. H. Immunoblot for ERα on RNA pulldowns using TFF1eRNA with increasing concentration of the TFF1 DNA as a competitor. I. TFF1eRNA RT-PCR from biotin labeled 3X ERE as bait with lysates from HEK-293T expressing either empty vector or ERα. J. Heatmap depicting the strength of ERα binding in intergenic regions within (fRIP interacting sites) and beyond (fRIP non-interacting sites) 10Kb of fRIP-seq peak.

The interactions between ERα and TFF1eRNA was validated and found to be strengthened upon ligand stimulation (Fig. 1F-G and Fig. S1C). Similar ligand-dependency was observed for various other RNAs (Fig. S1D). Next, we observed the binding of ERα with TFF1 eRNA was specific as addition of unlabeled DNA corresponding to TFF1 eRNA could not perturb this interaction (Fig. 1H). Further, TFF1 eRNA pulldown using biotin-3XERE (Estrogen Response Element) oligos as bait was enhanced upon ERα overexpression suggesting ERα can recruit TFF1 eRNA on ERE (Fig. 1I). Furthermore, we noted high ERα binding within 10 kb of fRIP-seq peaks (Fig. 1J). Lastly, this trend was more pronounced for the RNA in nucleoplasmic fraction as opposed to the chromatin (Fig. S1E-G). Together, this data suggest that ERα specifically interacts with a variety of RNAs. Further, the binding of ERα on genome is high near to RNA that interacts with ERα.

### RNA binding mutant of ERα shows loss of binding genome-wide

ERα interacts with RNA through its hinge region (255-272 aa) (Oksuz et al. 2023; Steiner et al. 2022; Xu et al. 2021) where amino acid stretch, RRGG (259-262) shows specific binding to RNA. Similar to this report, we created RNA binding deficient mutant of ERα (RBM-ERα) by substituting RRGG to AAAA (Fig. S2A). The nuclear localization (Fig. S2B) and DNA-binding was unaffected of RBM-ERα (Xu et al. 2021). As expected, RBM-ERα lacked the binding with TFF1eRNA as compared to the WT-ERα (Fig. 2A). Since, ligand strengthens ERα:RNA interactions (Fig. 1G and Fig. S1D), we hypothesized that lack of RNA binding may affect the chromatin binding of ERα in ligand dependent manner. Towards this, we performed chromatin fractionation on cells expressing RBM or WT-ERα upon E2 stimulation and found considerable reduction of RBM-ERα in chromatin fraction as compared to WT-ERα (Fig. 2B). To extend this observation genome-wide, we performed ChIP-seq of RBM and WT-ERα upon E2 exposure. We noted, decreased binding of RBM-ERα genome wide as compared to the WT-ERα (Fig. 2C and Fig. S2C). Loss of binding was observed for most of the E2 regulated genes including highly inducible genes like TFF1 and GREB1 (Fig. 2D and Fig. S2D). To rule out the loss of RBM-ERα binding in genome due to technical reasons, we performed paired-factor ChIP (pfChIP) of ERα and CTCF in the same reaction tube. CTCF was chosen due to its ERα independent binding in the genome (Holdinget al. 2018). We plotted the total read counts for CTCF peaks and observed unaffected CTCF binding in WT or RBM-ERα (Fig. 2E). However, RBM-ERα binding was observed to be poor (Fig. 2F). These results confirmed the specificity of the reduced genomic occupancy of ERα-RBM. Notably, the loss of RBM-ERα binding was proportional to the level of RNA being transcribed at a given genomic site (Fig. 2G). The lost peaks were majorly intergenic and intronic while more than 50% of retained sites were promoter suggesting, promoters were less susceptible to RNA-dependent loss of ERα (Fig. 2H, S2E). The intersection of these ChIP-seq and fRIP-seq revealed that the RNA from the lost sites interacts strongly with ERα as compared to the RNA from retained sites (Fig. S2F). Hence, lost sites had greater dependency on RNA for ERα recruitment as compared to retained sites.

**Figure 2.**
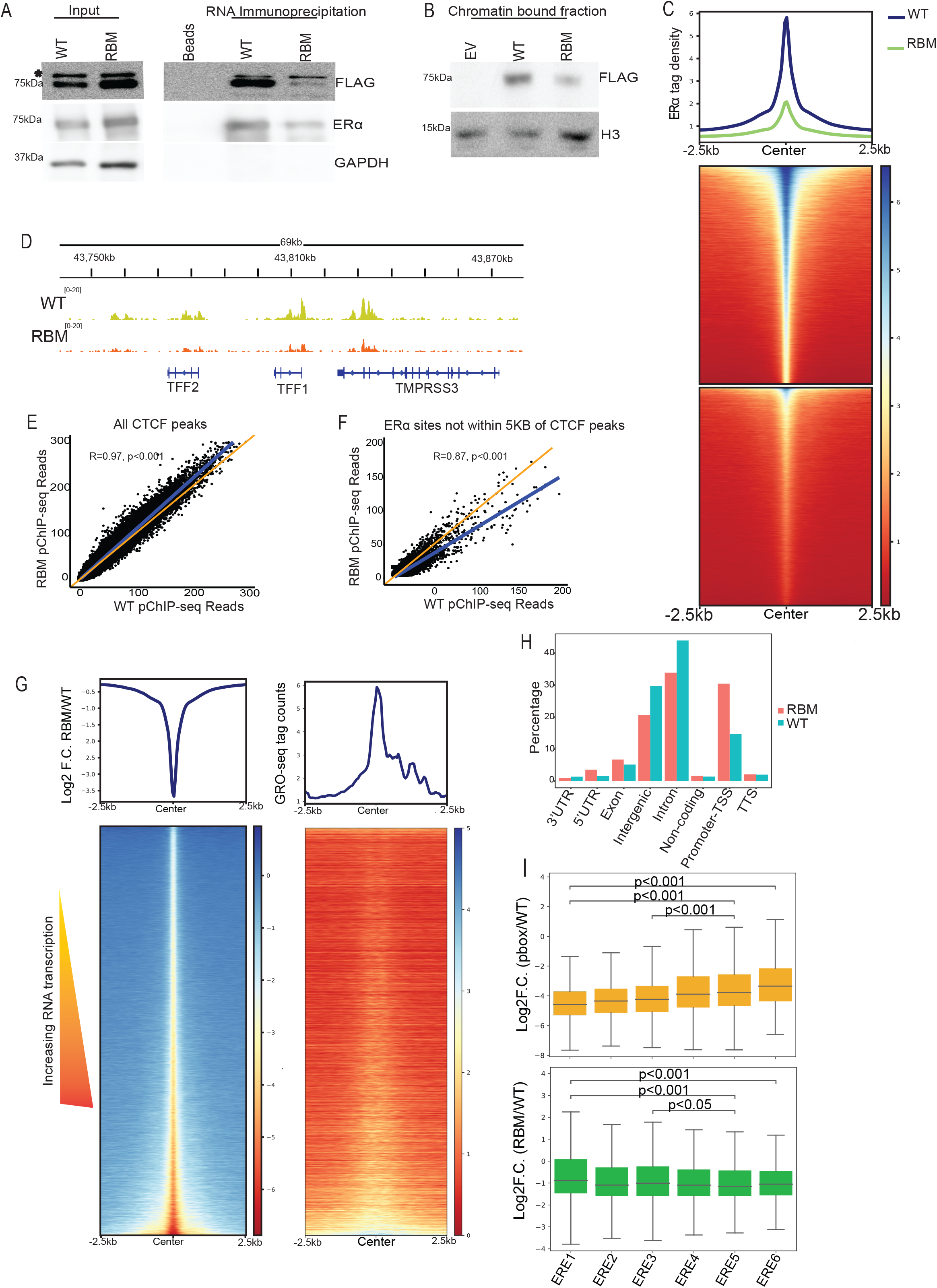
RNA binding mutant of ERα shows loss of binding genome-wide. A. Immunoblot for FLAG, ERα and GAPDH on RNA pull-downs using TFF1 eRNA for ERα WT and RBM over-expressed lysates from HEK-293T (* denotes non-specific band). B. Immunoblot for FLAG and H3 on chromatin bound fraction from ERα WT or RBM overexpressed MCF-7 cells. C. Heatmap depicting the binding strength of ERα:FLAG WT and RBM overexpressed in MCF- 7 cells. D. Genome browser screenshot for the binding of ER α WT and RBM on TFF1 locus in MCF-7. E-F. Normalized read count of ERα WT and RBM pfChIP-seq for all CTCF peaks and ERα peaks beyond 5kb of CTCF peaks respectively. G. Heatmap depicting the log2 F.C. of binding of RBM-ERα over WT-ERα and GRO-seq signal plotted at the sorted sites in the same order. H. Genomic distribution in percentages for the ERα:FLAG WT and RBM. I. Boxplot depicting the Log2F.C.of (pbox/WT) and (RBM/WT) at decreasing order of ERE strength. Statistical significance determined by Mann-Whitney U-test.

Additionally, binning of ERα peaks based on the strength of ERE revealed that the stronger ERE’s recruited ERα purely based on motif strength as these sites profoundly lost ERα, only upon mutation in DNA binding domain (DBD) or mutation in p-box (Fig. 2I, S3A and S3B). However, the lower strength ERE’s were dependent on RNA as these sites lost the most ERα, upon mutation in RBM but were least affected upon DBD perturbations (Fig. 2I, S3A and S3B). The lower strength ERE’s exhibited more RNA transcription and high relative H3K27ac (Fig. S3A). Furthermore depletion of total RNA using prolonged transcriptional inhibition followed by ERα ChIP-seq in T47D cells showed a similar pattern where sites with weak ERE’s lost more ERα as compared to the sites with stronger ERE’s (Fig. S3C) (Zhang et al. 2021). Together, these results suggest that recruitment of ERɑ by RNA is different from its DNA binding ability and the regions with weaker ERE’s depend on RNA for ERα recruitment.

### ERα Retention on Chromatin is RNA-Dependent

Next, we investigated whether high RNA-transcribing sites retain more ERα due to the RNA. To this end, we focused on non-genic ERα-bound sites from ChIP-seq because ERα exhibits the highest binding on these sites (Li et al. 2013). We grouped these sites into bins of nascent RNA tag counts from them in increasing order and observed a positive correlation between ERα binding and the level of transcription (Fig. 3A). Notably, this increase in ERα binding was independent of differences in ERE strength (Fig. 3B). Similar positive correlation between RNA and ERα was observed across all ERα-bound sites in the genome (Fig. S4A and S4B).

**Figure 3.**
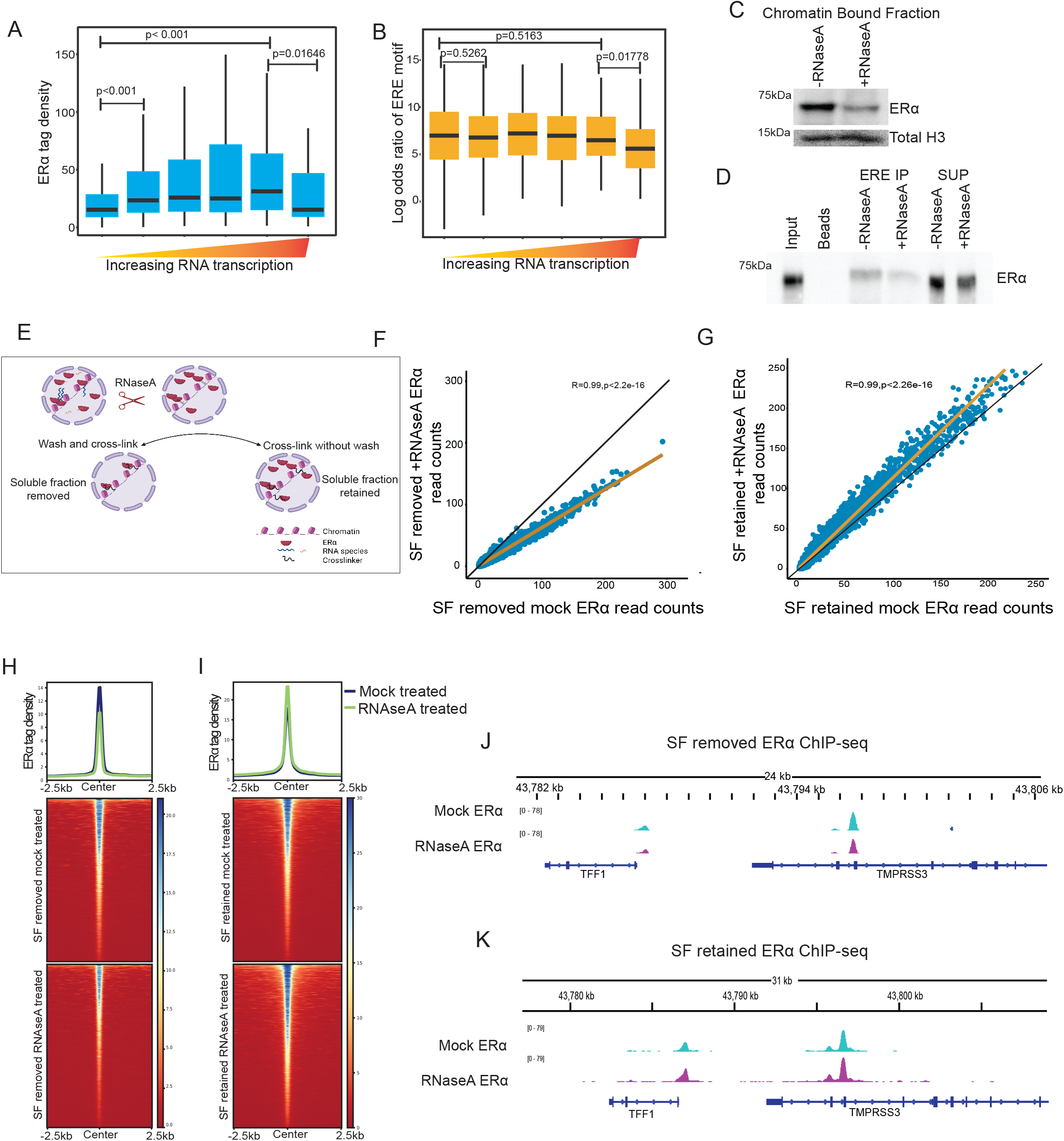
ERα retention on chromatin is RNA dependent. A. Boxplot showing ERα enrichment on all non-genic sites binned based on the levels of RNA transcription in increasing order. B. Boxplot showing the Log2 odds ratio for the ERE motif on all non-genic sites binned on the basis of transcription in increasing order. C. Immunoblot of ERα and total H3 on chromatin bound fractions from nucleus with or without RNase A treatment. D. ERE binding to ERα in cellular lysate in presence or absence of total RNA. E. Schematic of ChIP-seq following the removal of total RNA before crosslinking, with and without pre-extraction. F-G. Scatter plots showing correlation between ERα ChIP-seq normalized read counts with and without RNase A following pre-extraction and in the absence of pre-extraction respectively. H-I. Heatmap depicting ERα tag density between mock and RNase A treatment followed by pre-extraction and without pre-extraction respectively. J-K. Genome browser screenshot of TFF1 locus showing ERα ChIP-seq upon RNase A treatment with pre-extraction of soluble proteins and with retention of soluble proteins respectively.

To determine if ERα retention on DNA is dependent on RNA, we performed chromatin fractionation in the presence of RNAse A and found loss of ERα in chromatin fraction (Fig. 3C). Similar loss of ERα was also observed on EREs upon removal of total RNA (Fig. 3D). To extend these observations genome-wide, we performed RNase A treatment on non-crosslinked cells since crosslinking can compensate for the loss of binding (Thakur et al. 2019). We utilized a pre-extraction protocol following RNase A treatment to remove soluble and weakly bound chromatin fractions (Fig. 3E). The presence of this weakly bound subpopulation can yield contrasting results, as previously shown for CTCF (Gu et al. 2020; Saldaña-Meyer et al. 2019) and PRC2 (Beltran et al. 2016; Long et al. 2020). Upon performing ChIP-Seq after RNase A treatment and pre-extraction, we observed a severe loss of ERα binding in the genome (Fig. 3F, 3H, 3J and S4C). Conversely, ChIP-seq without pre-extraction showed excess binding of ERα in the genome (Fig. 3G, 3I, 3K and S4C). These findings suggest that RNA facilitates ERα binding on chromatin, and upon RNA removal, ERα interacts more but weakly with chromatin. This is seen as a loss of binding with pre-extraction and gain in binding without pre-extraction (Fig. S4D).

### RNA binding mutant of ERα interacts dynamically with the chromatin

The ChIP-seq results showed the reduced binding of RBM-ERα on its genomic sites (Fig. 2C and S2C) suggesting that RNA regulates the strength of ERα interaction with the chromatin. We hypothesized that the weak affinity of RBM-ERα could be associated with its dynamic molecular interactions in the nucleus including chromatin. To test this, we performed FRAP on WT:GFP and RBM-ERα:GFP in live cells (Fig. 4A). We observed that recovery of RBM-ERα after photo bleaching was much faster with median t ½ of 8 seconds as opposed to 12 seconds in the case of WT-ERα (Fig. 4B,C). The low affinity interactions can yield dynamic binding; we tested affinity of RBM-ERα binding to chromatin by fractionation of ERα from chromatin at varying salt concentrations. Indeed, the binding of mutant was of low affinity as seen by its elution from chromatin at low salt concentration (75-150 mM) as compared to the WT-ERα (300 mM) (Fig. 4D). Further, we confirmed the low affinity interactions of RBM-ERα on chromatin by retention assay where weakly chromatin associated proteins are removed from the nucleus using CSK (cytoskeleton) buffer. We noted loss of RBM-ERα over the WT-ERα suggesting weaker chromatin interactions (Fig. 4E). These results suggest that RNA binding is required for the stable binding of ERα to the chromatin as the loss of RNA binding results in more dynamic interaction of ERα with chromatin and higher nuclear mobility.

**Figure 4.**
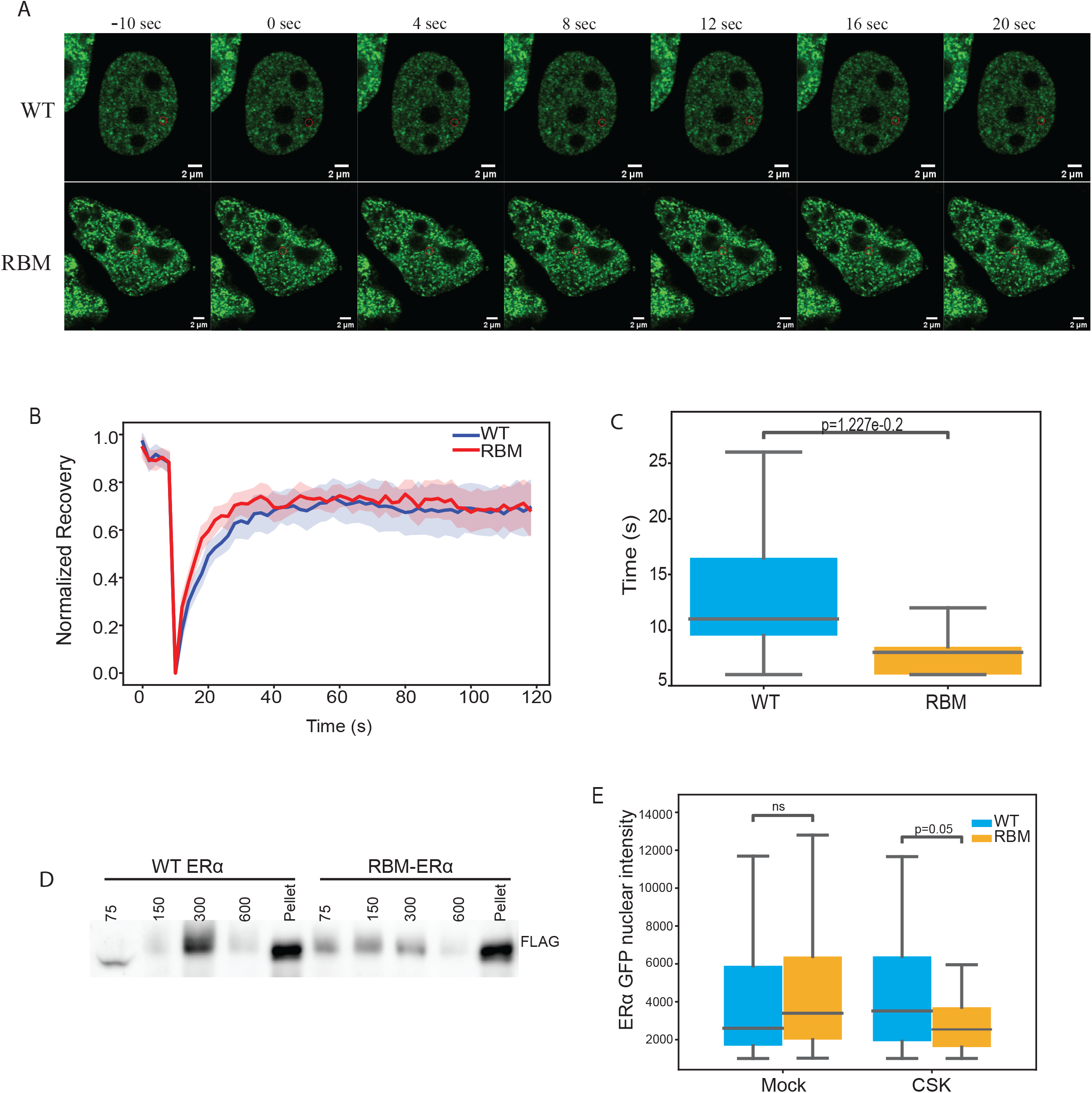
RNA binding deficit ERα interacts dynamically with the chromatin. A. Representative image of WT-ERα:GFP and RBM-ERα:GFP showing the FRAP ROI pre and post bleaching. B. Recovery plot of FRAP ROI for WT-ERα:GFP and RBM-ERα:GFP. C. Boxplot depicting the half-life recovery for WT-ERα:GFP and RBM-ERα:GFP. Statistical significance determined by unpaired t-test. D. Immunoblotting for FLAG on chromatin associated proteins eluted at different salt concentrations. E. Boxplot depicting the retention assay for WT-ERα:GFP and RBM-ERα:GFP. Statistical significance determined by Mann-Whitney U-test.

### The dynamic binding of ERα allows better transcription of target genes

To understand the impact of dynamic RBM-ERα binding on gene transcription, we conducted reporter assays using 3XERE upstream of the luciferase cassette in MCF-7 and HEK-293T cells. The RBM-ERα exhibited significantly higher luciferase activity compared to the WT-ERα (Fig. 5A, Fig.S5A-B). Expanding on this finding, we employed nascent EU-seq and observed a set of upregulated genes in cells expressing RBM-ERα over WT-ERα (Fig. 5B). Furthermore, we noticed a higher enrichment of PolIIS2p in the chromatin fraction of cells expressing RBM- ERα (Fig. 5C). To corroborate these observations, we performed total PolII ChIP-seq which also confirmed the greater occupancy of total PolII on the promoters of up-regulated genes identified from EU-seq (Fig. 5D). RBM-ERα had a more pronounced effect on the highly E2-induced genes (referred to as “changing” genes), as compared to less robustly E2-induced genes (referred to as “Non-changing”) (Fig. 5E-F). The changing genes also exhibited higher PolII occupancy compared to the non-changing and random genes upon RBM-ERα expression (Fig. 5G and 5I). The extent of PolII gain on the gene body of these changing genes was similar to the gain observed upon E2 treatment compared to vehicle (Fig. 5H).

**Figure 5.**
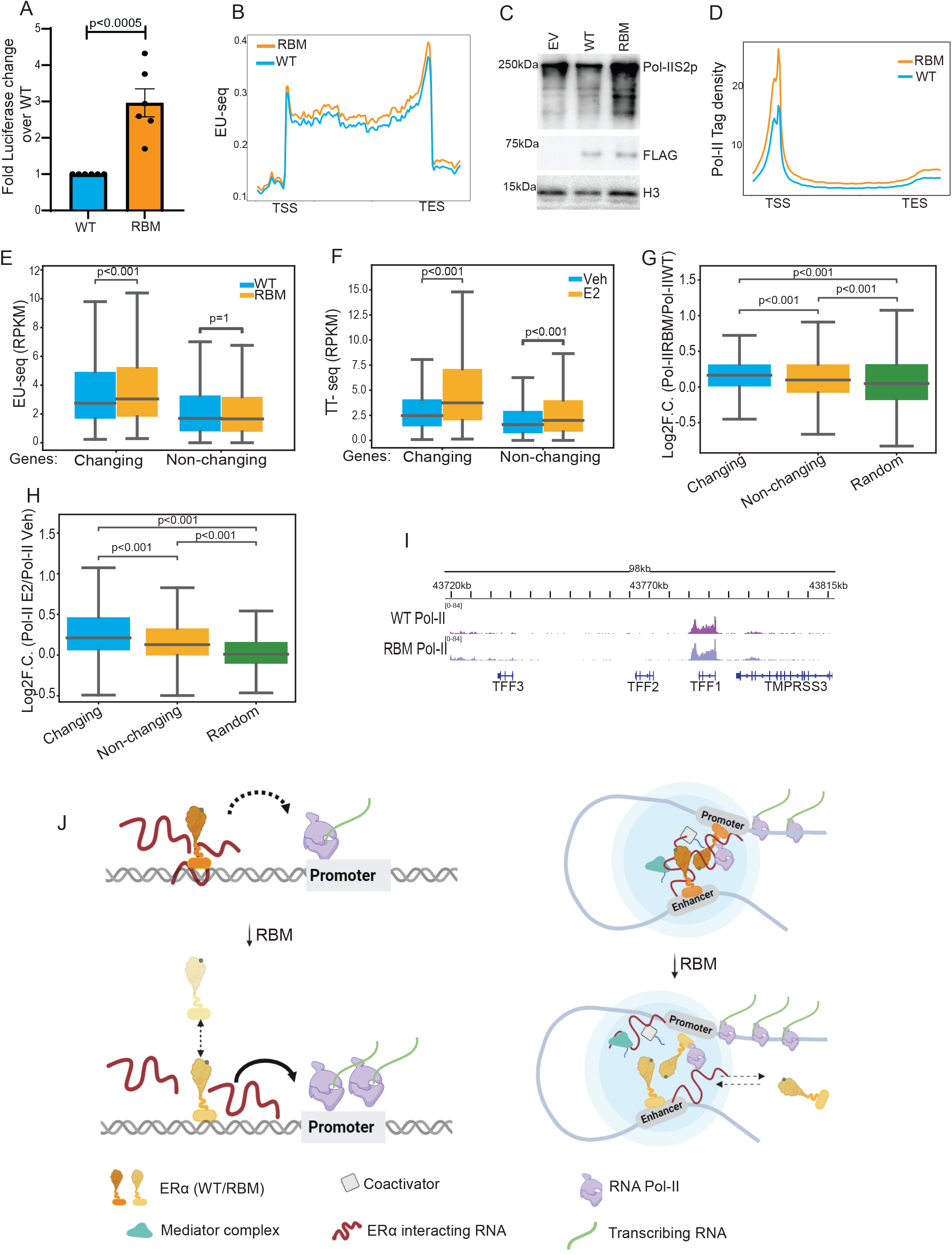
The dynamic binding of ERα allows better transcription of target genes. A. Luciferase activity normalized to the WT for 3X ERE reporter assay with ERα WT and RBM overexpression in MCF-7 upon 4 hours of E2 treatment. Statistical significance determined by Mann-Whitney U-test. B. Summary plot showing the EU-seq reads from whole gene body for up-regulated genes upon RBM-ERα expression over WT-ERα. C. Immunoblot of PolIIS2p, FLAG and H3 from chromatin bound fractions of either empty vector or WT-ERα or RBM-ERα expressing nuclei. D. Summary plot showing the total PolII ChIP-seq reads from whole gene body for upregulated genes upon RBM-ERα expression over WT-ERα. E. Boxplot showing the EU-seq reads for the whole gene body in two categories- changing genes and non-changing genes in WT-ERα or RBM-ERα. Statistical significance determined by Wilcoxon signed rank sum test. F. Boxplot showing the TT-seq reads for the whole gene body upon vehicle or E2 treatment in two categories- changing genes and non-changing genes. Statistical significance determined by Wilcoxon signed rank sum test. G. Boxplot depicting the log2F.C.for PolII ChIP-seq (RBM-ERα/WT-ERα) for three categories namely, changing genes, non-changing genes and random gene list. Statistical significance determined by Mann-Whitney U-test. H. Boxplot depicting the Log2F.C. of PolII ChIP-seq(E2/Veh) for three categories namely, changing genes, non-changing genes and random gene list. Statistical significance determined by Mann-Whitney U-test. I. Genome browser screenshot of TFF1 locus for the total Pol-II ChIP-seq upon ERα WT or RBM overexpression. J. Model

Additionally, we found that a substantial proportion of ERα binds to introns at both DNA and RNA levels (Fig. S5D-E and Fig. 1A). Notably, the changing genes displayed more intronic RNA-ERα interactions and more ERα binding on the gene body, resulting in the greater loss of ERα upon RBM-ERα mutation (Fig. S5F-H). These results suggest that lack of RNA binding in case of RBM-ERα allows it to bind on target sites in dynamic manner which favors the robust PolII recruitment and transcription.

## Discussion

### ERα interacts with RNA

The interaction of TFs with DNA was assumed to be the canonical mode of TF binding on chromatin. Recent studies dissecting the functional role of TF-RNA interactions have revealed that RNA modulates the TF’s interaction with chromatin leading to transcription regulation. The interaction between ERα and RNA has been shown (Oksuz et al. 2023; Steiner et al. 2022; Xu et al. 2021; Yang et al. 2020). However, its role in ERα binding on chromatin and downstream effects on transcription is controversial from presumed no effect on DNA binding and transcription under basal signaling (Xu et al. 2021) to significant effect on transcriptional activation (Oksuz et al. 2023). The ligand modulates the binding of ERα on DNA and facilitates the recruitment of robust transcription machinery for target gene activation (Shang et al. 2000) suggesting, ERα:RNA interaction and its function may be influenced by the ligand-dependent events. Therefore, we utilized E2 signaling to examine the nuclear role of ERα:RNA interaction as opposed to basal signaling (Xu et al. 2021). Our findings reveal that ERα binds to various coding and non-coding RNAs (Fig. 1A), and this interaction is enhanced upon E2 stimulation (Fig. 1G and S1D). Specifically, we confirm that ERα interacts with RNA through the hinge region in c-terminal extension (CTE) of its DNA-binding domain (DBD) (Fig. 2A, 1H-I). This RNA-dependent mechanism of ERα recruitment differs from previous findings (Yang et al. 2020) where repressive eRNAs interacted with ERα solely through the DBD thus, precluding simultaneous interactions with both DNA and RNA. Our findings align with earlier studies on GR and Sox2, demonstrating that these transcription factors can interact with specific RNA either via their DBD or the CTE of DBD (Holmes et al. 2020; Hou et al. 2020; Lammer et al. 2023; Parsonnet et al. 2019), thereby modulating their DNA occupancy. We provide evidence supporting the functional implications of ERα’s concurrent DNA-RNA binding whereby, RNA interactions with ERα result in higher occupancy compared to non-interacting sites (Fig. 1J). The binding of ERα to RNA offers an additional mechanism for its chromatin tethering while its DBD engages with ERE on DNA.

### RNA supports ERα binding on weaker ERE motifs

Owing to ERα’s capability to bind RNA, our data suggest that transcribed sites with similar ERE strength can recruit more ERα as compared to non-transcribing sites (Fig. 3A-B). Furthermore, pronounced loss of RBM-ERα on weak EREs, as opposed to the loss of DBD mutant ERα on strong EREs (Fig. 2I and S3A) confirms the RNA mediated binding of ERα on weaker EREs. Intriguingly, though the ERα binding is highest at the strong EREs, the weaker or low-affinity EREs display higher levels of transcription and H3K27ac activation marks suggesting that low-affinity binding sites play a significant role in driving robust transcription.

In addition to the major groove interactions mediated by the DBD, the CTE of nuclear receptors including ERα establishes additional contact with the minor groove of DNA (Jakób et al. 2007; Gearhart et al. 2005; Zhao et al. 1998). Based on these facts, we propose that the interaction of RNA with the CTE of ERα, can stabilize ERα on DNA. Moreover, in the absence of RNA or upon RBM mutation, ERα loses its ability to contact RNA that leads to destabilization of ERα on chromatin and results in its dynamic binding behavior. These observations were supported by retention assays, salt elution and FRAP (Fig. 4 and Fig. 3). This increased mobility may arise from the loss of RNA-mediated entrapment, as RNA can act as a crosslinker. The absence of this crosslinking effect may allow ERα to be more available for chromatin binding in dynamic manner. Indeed, the inability of TFs to interact with RNA increases its unbound fraction in the nucleus as revealed by single particle tracking (Oksuz et al. 2023).

### Dynamic binding of TF with chromatin corelates with robust transcription

As a result of polymerase loading, the transcription takes place in bursts as opposed to constant transcription (Rodriguez et al. 2019, Pomp et al. 2023). The occurrence of bursts is anti-correlated with binding of transcription factors (Charoensawan, Martinho, and Wigge 2015; Doidy et al. 2016; Para et al. 2014; Pownall et al. 2023; Schaffner 1988). Therefore, the high burst frequencies require not the static binding of TF but a dynamic binding with each binding event long enough (dwell time) to result in subsequent transcription burst. Indeed, it is now established that TF binding is much more dynamic as opposed to long prevailed static binding model (Hager, McNally, and Misteli 2009; Hansen et al. 2019; Swinstead et al. 2016) and further, stable binding is inhibitory to transcription (de Jonge et al. 2020).

In this direction, enhanced mobility of ERα-RBM potentially results in the higher PolII occupancy and transcription of E2-regulated genes (Fig. 4C-E and Fig. 5). Each dynamic binding event of mutant ERα might be long enough to engage in PolII loading. Further, these observations are consistent with previous reports showing that the dynamic binding of ERα and AR to chromatin lead to its enhanced transcriptional activity (Guan et al. 2019; Kim et al. 2022). Similarly in the case of AR, mutations in the hinge region have been reported to cause increased transcription and mobility inside the nucleus (Buchanan et al. 2001; Haelens et al. 2007; Tanner et al. 2010).

### ERα:RNA behavior under basal signaling vs. post ligand stimulation

Under basal signaling, unliganded ERα poorly binds to chromatin and transcription activating machinery (Shang et al. 2000) and, preferentially interacts at 3’UTRs of mRNAs to regulate the integrated stress response through their translation regulation (Xu et al. 2021). Understandably, under basal signaling, Xu et al., did not observe the loss of RBM-ERα binding on the genome and therefore, the effect on transcription. Similar to Xu et al., our study also focuses on RRGG motif in hinge domain of ERα but under ligand stimulation.

Similar to our and Xu et al., 2021, Oksuz et al., 2023 have also revealed the interaction of ERα with RNA through its arginine-rich motif (ARM) specifically, at R269 in the same hinge domain (255-272 aa). Further, similar to our observation, RBM of ERα (R269C) also shows loss of binding to the RNA and chromatin. However, Oksuz et al. show reduced trans-activation potential of ARM-ERα (R269C) using Tat-reporter assays. This assay was performed by using ERα-ARM and ARM-ERα (R269C) peptide truncation without DNA binding domain of ERα. Tat- reporter assays require binding of ERα-ARM to RNA for transcriptional activation of reporter whereas, we have used EREs (DNA) that rely on ERα binding through its DBD for reporter gene activation. The DBD based reporter of R269C also show activation of reporter activity as shown previously (Boldes et al. 2020). These data suggest that the role of TF’s RNA binding along with its DNA binding ability might be different from its trans-activation potential in absence of DNA binding. Our PolII ChIP-seq and EU-seq also support the results from reporter assays. Further, the mutation of different hinge residues in ours and Oksuz et al. may result in varying transcriptional responses (Haelens et al. 2007) for example, 269R in Oksuz et. al. and RRGG in our study may have different biological functions owing to RNA specificity. Future studies on these differences will be helpful.

## Author’s Contribution

The study was conceptualized and designed by DS and DN. Most experiments were performed by DS with help from RM and TK. DS and BS performed NGS data analysis. The manuscript was written by DS and DN. All authors read and edited the manuscript.

## Acknowledgments

We acknowledge support of the Department of Atomic Energy, Government of India, under project no.12-R&D-TFR-5.04-0800 and intramural funds from NCBS-TIFR (to DN). DN is an EMBO Global Investigator. We also acknowledge the funding support from Welcome-IA (IA/1/14/2/501539) and DST core grant (CRG/2019/005714) to DN. DS, TK and RM are supported by the TIFR-NCBS graduate program. We acknowledge the support from core facility at NCBS specifically NGS and CIFF. DS acknowledges the help from Ananya Sadhu, Awadhesh Pandit, Kuldeep Sachdeva and Arif Hussain for help in experiments. We thank Sachin Mishra and Amanjot Singh for critical comments on the manuscript. We thank DN lab members for discussions.

## Material and Methods

### Cell culture

MCF-7 and HEK-293T cells were obtained from ATCC and maintained in high glucose DMEM (Invitrogen) at 37℃ with 5% CO2. For hormone deprivation, MCF-7 cells were seeded in complete DMEM and the following day, cells were washed with 1X DPBS and then cultured in DMEM without phenol red (Invitrogen) supplemented with 5%charcoal-stripped FBS (Invitrogen). After 72 hours of hormone deprivation, the cells were treated with either 100nM 17β estradiol (Sigma E2758) or vehicle (ethanol, Millipore) for 1 hour for all experiments. For the over-expression studies, transfection was performed using Lipofectamine 2000 (Invitrogen 11668019) according to the manufacturer’s protocol in OptiMEM with reduced serum and no phenol red (Invitrogen 11058021) after48 hours of hormone deprivation. After 6 hours of transfection, the media was changed to stripping media, and on the third day, the cells were stimulated with the ligand.

### Formaldehyde Assisted RNA Immunoprecipitation (fRIP)

The fRIP procedure was performed following the previously described protocol (Hendrickson et al. 2016). Briefly, MCF-7 cells were crosslinked with 0.1% formaldehyde for 10 minutes at room temperature (RT) with gentle shaking. The crosslinking reaction was quenched by the addition of 0.125M glycine for 5 minutes at RT. The cells were then washed, pelleted, and stored at - 80°C. For cell lysis, approximately 20 million cells were resuspended in 1250μl of RIPA lysis buffer (50mM Tris pH 8, 150mM KCl, 0.1% SDS, 1% Triton-X, 5mM EDTA, 0.5% sodium deoxycholate, 0.5mM DTT, 1X protease inhibitor cocktail (PIC), 100 U/ml RNase inhibitor) and incubated on ice for 10 minutes. Then the lysates were sonicated using a Bioruptor Pico (from Diagenode) for 6 cycles with 30 seconds ON and 30 seconds OFF program. The lysates were then clarified by centrifugation at 15,000 rpm for 10 minutes, and the supernatant was diluted by adding an equal amount of fRIP wash buffer (150mM KCl, 25mM Tris pH 7.5, 5mM EDTA, 0.5% NP-40, 0.5mM DTT, 1X PIC, 100 U/ml RNase inhibitor). The diluted lysates were passed through a 0.45μM membrane syringe filter. For immunoprecipitation, 1μg of anti-ERα antibody (sc-8002) was added to 1 ml of lysate, followed by incubation at 4°C for 2 hours. After incubation, 20μl of Protein G beads were added, and the mixture was incubated for 1 hour on a rotor at 4°C. The antibody-protein-bead complexes were then washed thrice with fRIP wash buffer at high speed on the rotor.

### For RNA elution, purification, and fRIP-PCR/fRIP-seq

The beads were resuspended in 56μl of nuclease-free water (NFW). Then, 33μl of elution buffer was added to each tube. The samples were incubated at 42°C for 1 hour, followed by an additional incubation at 55°C for 1 hour. TRIzol was added to the tube to extract RNA according to the manufacturer’s instructions. The purified RNA was used for cDNA synthesis using the SuperScript™ IV First-Strand Synthesis System (#18091050) or for preparing RNA-seq libraries, using the NEBNext rRNA Depletion Kit v2 and NEBNext Ultra II Directional RNA Library Prep, following the manufacturer’s protocol.

### fRIP-seq analysis

The adaptor sequences were removed from the reads using Cutadapt (Martin 2011), followed by alignment to the hg19 reference genome using RNA STAR (Dobin et al. 2013). Duplicate reads were removed using UMI-tools deduplicate (Smith, Heger, and Sudbery 2017). Regions of fRIP-seq IP enrichment were called using MACS2 (Feng et al. 2012; Zhang et al. 2008) with the fRIP-seq input as the control file. The log2FC of fRIP-seq IP /fRIP-seq input was calculated using the aligned BAM file with bamcompare (Ramírez et al. 2016). For visualization purposes, read densities were calculated across the genome as bedgraph files and uploaded onto the IGV genome browser.

### RNA immunoprecipitation Assay

The genomic region transcribing TFF1 eRNA was amplified by PCR using the primers listed in Supplementary Table S1. The amplified PCR product was then cloned into the pcDNA3.1+ vector downstream of the T7 promoter. The digested vector was subjected to invitro transcription (IVT) following the manufacturer’s protocol using T7 polymerase (Promega P2075) and biotin RNA labeling mix (Roche 11685597910).

RIP was performed as previously described with minor modifications (Jayani, Singh and Notani 2017). Briefly, MCF-7 cells were washed and pelleted using 1X PBS. For cell lysis, 1ml of RIP buffer (25mM Tris-HCl pH 7.4, 150mM KCl, 0.5mM DTT, 0.5% NP-40, 1X protease inhibitor cocktail, and 100 U/ml RNase inhibitor) was added to 5 million cells, followed by 5 minutes of incubation on ice and 10 cycles of sonication using a Bioruptor pico with 30 seconds ON and 30 seconds OFF settings. The lysates were then clarified by centrifugation at 12,000 rpm for 12 minutes. To the cell lysate, 2μg of IVT-biotin labeled TFF1 eRNA fragments were added to each tube and incubated on the rotor for 2 hours at 4°C. 15μl of Dynabeads MyOnestreptavidin beads were added and incubated for an additional 45 minutes. The beads were washed twice with RIP buffer containing 300mM KCl, and the proteins were eluted by boiling the beads at 98°C in 2X lamellae dye.

For the DNA competition RIP assay, unlabeled purified PCR product of the same genomic region was added to the cell lysates in three different conditions: No DNA, 1XDNA concentration and 2XDNA concentration. For the WT and RBM-ERα RIP assay, ERα WT or RBM constructs were transfected into HEK-293T cells. After 24 hours of transfection, the cells were treated with 100nM E2 and harvested to perform RNA pulldowns.

### 3X ERE Biotin pulldowns

To perform ERE pulldowns, 3XEREoligoand its reverse complementary sequence (provided in Table S2) were annealed, and end labeling was performed using T4 PNK from NEB (cat number: M0201S) and biotin-14-dATP from Invitrogen (cat number: 19524016).

For the ERE pulldowns, cell lysates were either mock-treated or treated with RNase A (Qiagen 19101) and then incubated at 4°C for 2 hours with biotinylated oligos. After 2 hours, 20μl of DynabeadsMyOne streptavidin beads were added and incubated for another 45 minutes. The beads were then washed three times with NP40 lysis buffer, and the proteins were eluted by boiling at 98°C in 2X Laemmli dye.

### For the TFF1 eRNA capturing experiment

HEK-293T cells were transfected using Lipofectamine 2000 according to the manufacturer’s protocol with either the empty vector or a vector over-expressing ERα. Cells were then ligand stimulated, and nuclei isolation was performed as previously described(Liu et al. 2014).Briefly, the cell pellet was resuspended in 1ml of hypotonic buffer (10mM Tris-HCl pH 7.5, 2mM MgCl2, 3mM CaCl2, and 1X protease inhibitor cocktail) and incubated on ice for 5 minutes, followed by centrifugation at 3000 rpm for 5 minutes at 4°C. The swollen cell pellets were lysed by resuspending in lysis buffer (hypotonic buffer + 0.5% NP-40 + 1X protease inhibitor cocktail) and incubated on ice for 5 minutes. The nuclei were collected by centrifugation at 6000 rpm for 10 minutes. The nuclei pellet was then resuspended in NP-40 lysis buffer (50mM Tris-HCl pH 7.4, 150mM NaCl, 1% NP-40, 0.5% sodium deoxycholate, 0.1% Triton X-100, and 1X protease inhibitor cocktail), incubated on ice for 10 minutes, and clarified by centrifugation at 12,000 rpm for 12 minutes. To the 500μl of nuclear lysates prepared from either the empty vector or ERα over-expressed HEK-293T cells, 0.2μM of the biotin-labeled3XERE oligos were added (sequence listed in Table S2). Unlabelled IVT TFF1 eRNA (Fragment1) was sequentially added to each tube, and the reaction was incubated on a rotor at 4°C for 2 hours. Then, 25μl of DynabeadsMyOne streptavidin beads were added and incubated for another 45 minutes. The beads were washed three times with NP40 lysis buffer, and 1ml of TRIzol (Invitrogen 15596026) was added to the beads to isolate the RNA. cDNA was prepared, and PCRs were performed using TFF1eRNATFF1 eRNA

### Subcellular fractionation

To isolate cytoplasmic, nucleoplasmic, and chromatin-bound proteins, the method described (Caudron-Herger et al. 2019) was adopted. Briefly, cells were swelled and lysed in hypotonic buffer (10mM Tris HCl at pH 7.5, 10mM NaCl, 3mM MgCl2, 0.3% NP-40, 10% glycerol, 1X protease inhibitor cocktail) by incubating on ice for 10 minutes. The lysate was then centrifuged at 3000 rpm for 5 minutes, and the supernatant was transferred to another tube and labelled as the cytoplasmic fraction. The isolated nuclei were washed three times with HLB buffer and then resuspended in modified Wuarin-Schiebler buffer (MWS; 10mM Tris-HCl at pH 7.5, 300 mM NaCl, 4mM EDTA, 1M urea, 1% NP-40, 1% glycerol, 1X protease inhibitor cocktail). For RNAseA treatment of the nucleoplasmic fraction, it was divided into two equal parts, with one part receiving RNAseA at a concentration of 2μg/ml, while the other part served as the control. The nucleoplasmic fractions were incubated on ice for 10 minutes. After incubation, the samples were spun at 5000 rpm for 5 minutes, and the supernatant was labelled as the nucleoplasmic fraction. To isolate the chromatin-bound fraction (CBF), nuclear lysis buffer (NLB: 20mM Tris HCl at pH 7.5, 150mM KCl, 3mM MgCl2, 0.3% NP-40, 10% glycerol, 1X protease inhibitor cocktail) was added, followed by sonication for 10 cycles with 30 seconds ON and 30 seconds OFF settings. To all the fractions, SDS dye was added, and the samples were boiled at 98°C for 10 minutes.

### ChIP-seq

MCF-7 cells were seeded and hormone-stripped for 48 hours. After that, the cells were transfected with either the WT ERα or RBM-ERα expressing plasmids. Following 24 hours of transfection, the cells were treated with 100nM E2 for 1 hour and crosslinked with 1% formaldehyde for 10 minutes at room temperature with gentle shaking. The crosslinking reaction was quenched by adding 0.125 M glycine for 5 minutes at room temperature. ChIP-seq samples were prepared according to the protocol described (Saravanan et al. 2020).

Briefly, the cells were washed thrice with 1X PBS and scraped in 1X PBS. The cell pellets were then spun at 3000 rpm for 5 minutes to pellet the cells, and the pellets were stored at -80°C until further use. Approximately 10 million cells were resuspended in 1 ml of nuclear lysis buffer (NLB) (50mM Tris-HCl pH 7.4, 1% SDS, 10mM EDTA pH 8.0, and 1X PIC) and incubated on ice for 10 minutes. After incubation, the samples were sonicated using a Bioruptorpico for 28 cycles with 30 seconds ON and 30 seconds OFF settings. The lysates were then clarified by centrifugation at 12,000 rpm for 12 minutes. A total of 100 μg of chromatin was measured and diluted 2.5 times with dilution buffer (20mM Tris-HCl pH 7.4, 100mM NaCl, 2mM EDTA pH 8.0, 0.5% Triton X-10, and 1X PIC). A total volume of 500 μl was used for each IP. Additionally, 50 μl was taken out and labeled as 10% input. For each IP sample, 1 μg of either anti-FLAG (Sigma F7425) or anti-PolII (Diagenode C15200004 and sc-55492) antibody was added and incubated overnight at 4°C on a rotor. For parallel ChIP-seq, both anti-FLAG (Sigma F7425) and anti-CTCF (CST 3418S) antibodies were added simultaneously in the same tube. To pull down the antibody-protein DNA complex, 15 μl of Protein G beads were added the next day and incubated for 4 hours at 4°C on the rotor. After incubation, the beads were washed sequentially with wash buffer I, wash buffer II, wash buffer III, and 1X TE. The DNA-protein complexes were eluted from the beads by resuspending them in 200 μl of elution buffer (100mM NaHCO3, 1% SDS) at 37°C in a thermomixer with 1400 rpm. The eluted DNA-protein complexes were reverse-crosslinked by the addition of 14 μl of 5M NaCl and overnight incubation at 65°C. The ChIP DNA was purified using phenol:chloroform:isoamyl alcohol and ethanol precipitation. The resulting pellet was resuspended in 10 μl of nuclease-free water (NFW) and used for ChIP-seq library preparation using the NEBNext® Ultra™II DNA Library Prep kit with Sample Purification Beads (E7103L) following the manufacturer’s protocol.

### ChIP-seq analysis

The raw reads obtained from the sequencing were aligned to the hg19 reference genome using Bowtie2 (Langmead and Salzberg 2012). Reads with a quality score less than 20 were filtered out using the Filter Sam or Bam tool (Li et al. 2009). Peaks were called using MACS2 (Feng et al. 2012) with default settings. The resulting peaks were visualized using the IGV genome browser. For pfChIP-seq, the total read counts were plotted for all CTCF peaks and ERα peaks that are not within a 5kb distance of CTCF peaks.

### RNase A ChIP-seq

MCF-7 cells were subjected to E2 stimulation, washed with 1X PBS three times, and pelleted. RNase A ChIP-seq was performed following a previously described protocol (Thakur et al. 2019; Zoabi et al. 2014) with slight modifications. Briefly, the cell pellet was resuspended in a buffer (20mM HEPES pH 7.5, 0.1mM CaCl2, 3mM MgCl2, 150mM NaCl, 0.05% NP-40, and 1X PIC). The cell suspension was divided into two equal parts. One aliquot was treated with RNase A at a concentration of 2µg/ml, while the other aliquot was mock treated. Both aliquots were incubated on ice for 10 minutes. For the ChIP-seq with pre-extraction, the cells were pelleted by centrifugation at 3000 rpm at 4°C and washed with the resuspension buffer. The cell pellets were then resuspended in 500 µl of resuspension buffer and crosslinked using 1% formaldehyde for 10 minutes at room temperature. The crosslinking reaction was quenched with 0.125 mM glycine. For the ChIP-seq without pre-extraction, the cells pellet was resuspended in resuspension buffer (20mM HEPES pH 7.5, 0.1mM CaCl2, 3mM MgCl2, 150mM NaCl, 0.05% NP-40, and 1X PIC). The suspension was divided into two equal parts, one aliquot was mock treated and other was treated with RNAse A for 10 minutes on ice. After incubation time 1% formaldehyde was directly added to the cell suspension, followed by quenching with 0.125mM glycine.

### Generation of ERα RBM mutant plasmids

The WT ERα and RNA binding mutants (259-262 RRGG>AAAA) constructs were generated by PCR amplification using the oligos listed in tableS3. The amplified products of the WT and RBD mutants were then cloned into the 3XFLAG CMV10 vector at the EcoRI and BamHI restriction sites.

### Gradient salt elution for chromatin bound proteins

Nuclei were isolated from cells expressing either wild-type (WT) or RBM FLAG-tagged ERα following the protocol mentioned above. The isolated nuclei were then resuspended in a buffer containing 10mM Tris HCl (pH 7.5), 0.15% NP-40, 2mM EDTA, 1X PIC and 75mM NaCl. The nuclei suspension was rocked at 4℃ for 15 minutes to allow for protein extraction. After incubation, the nuclei were pelleted by centrifugation at 4000rpm for 5 minutes, and the supernatant was collected. This elution step was repeated multiple times, each time using an increasing NaCl concentration. Finally, after the 600mM NaCl elution, the pellet was resuspended in 2X lamellae dye, and the samples were boiled at 98℃ for 10 minutes. This process allows for the extraction and collection of proteins associated with the chromatin-bound fraction.

### Retention Assay

Cells were grown on coverslips and transfected with ERα:GFP WT or RBM after 48 hours of hormone deprivation. After 24 hours of transfection, cells were stimulated with E2 for 1 hour. The coverslips were washed three times with 1X PBS. For the pre-extraction of weakly associated chromatin-bound proteins, cells were treated with CSK buffer (10mM PIPES/KOH pH 6.8, 100mM NaCl, 300mM sucrose, 1mM EGTA, 1mM MgCl2, 1mM DTT, and 1X PIC) for 5 minutes on ice. Another set of coverslips were mock treated to serve as a control. After the incubation, the CSK buffer was removed, and cells were crosslinked using 4% paraformaldehyde. The coverslips were mounted with 90% glycerol. Cell imaging was performed using a 60x/1.42 oil objective on the Olympus FV3000 microscope.

### Dual Luciferase Assay

MCF-7 cells were grown in a 24-well plate using hormone-deprived media for 48 hours. Subsequently, the cells were co-transfected with 250ng of 3X ERE with TATA-box luciferase plasmid (Addgene 11354), a gift from Donald McDonnell, 2.5ng of Renilla TK plasmid, and either WT or RBM-ERɑ. The cells were then treated with E2 for either 24 hours or 4 hours. Luciferase activity was measured using the Multimode Reader Varioskan Lux, with Renilla luciferase used as an internal control. Multiple biological replicates were performed without technical replicates.

### ERE binning

ERɑ peaks were called using MACS2 (Feng et al. 2012) and motif scores were calculated using FIMO (Grant, Bailey, and Noble 2011). Out of a total of 11,763 peaks, 5,311 regions showed the presence of the ERE motif. These regions were divided into six continuous bins based on the decreasing order of the ERE motif score. ERɑ tag density, Log2F.C. (ΔDBD/WT), Log2F.C. (pBox/WT), Log2F.C. (RBM/WT), TT-seq RPKMs, and H3K27ac tag density were plotted for these regions.

### EU-seq

MCF-7 cells were grown in hormone-deprived media for 48 hours and transfected with either the WT-ERα or RBM-ERɑ. After 6 hours, the media was changed. Following 24 hours of transfection, the cells were treated with 100nM of E2 for 1 hour, during which nascent RNA was labeled with 0.5mM EU for 45 minutes. Total RNA was isolated using TRIzol (Invitrogen 15596026) according to the manufacturer’s protocol. rRNA depletion was performed using the NEBNext® rRNA Depletion Kit v2 (E7405L). The nascent EU-labeled RNA was biotinylated using biotin azide and captured using DynabeadsMyOne Streptavidin T1 magnetic beads, following the manufacturer’s protocol (Click-iT™ Nascent RNA Capture Kit, for gene expression analysis, C10365). Subsequently, cDNA was synthesized using the SuperScript™ IV First- Strand Synthesis System (18091050). The second strand of cDNA was synthesized using the NEBNext® Ultra™ II Directional RNA Library Prep with Sample Purification Beads (Catalogue no-E7765L). The double-stranded DNA was further fragmented, and the library was prepared using the NEBNext® Ultra™ II FS DNA Library Prep Kit for Illumina (E7805L), following the manufacturer’s protocol.

### EU-seq analysis

The raw reads were aligned to hg19 using Bowtie2 (Langmead and Salzberg 2012). PCR duplicates were removed using MarkDuplicates from Picard Tools (Anon n.d.). Read coverage on known E2 upregulated genes was calculated using multiBamSummary (Ramírez et al. 2016) and the gene list was categorized into changing and unchanging categories.

### FRAP

The FRAP assay was performed after 45 minutes of E2 (estradiol) treatment using a 488nm laser. Transfected cells were bleached with 100% laser power over an area of 1μm, and images were collected every two seconds after bleaching. Prior to bleaching, five frames were collected. The bleaching process was conducted in the sixth frame, and images were taken for the following two minutes. The intensity of the Region of Interest (ROI) was measured across the time frames. To account for cell-to-cell variability and ROI differences, all the frames were normalized using the mean intensity of the ROI in the first frame. Additionally, the curve was min-max normalized to correct for bleaching differences. For half time calculation the time it takes for the normalized intensity to recover to half of its original value was calculated.

---

**Supplementary Figure S1.**
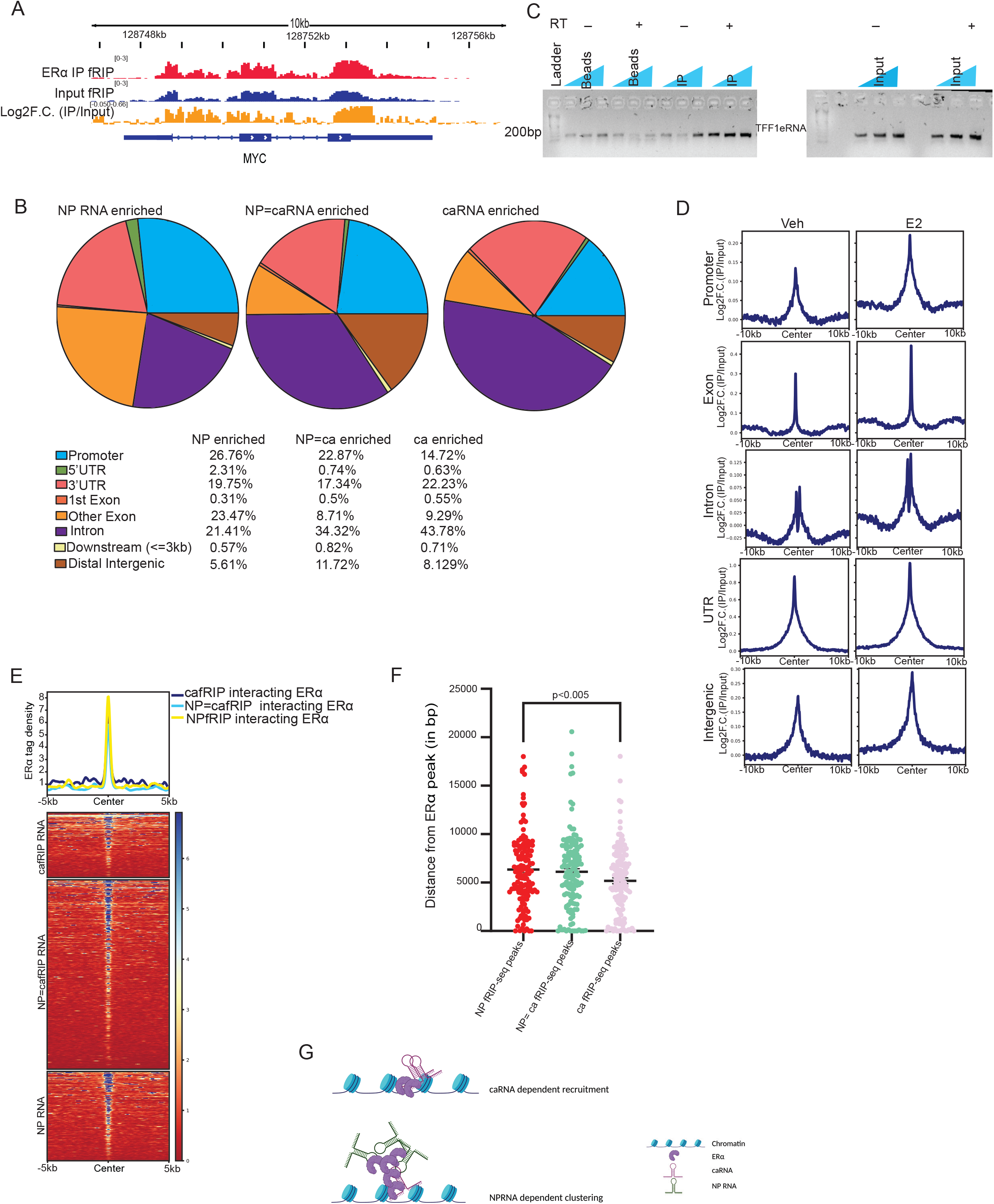
(Related to Figure 1) A. Genome browser screenshot showing fRIP-seq IP, Input and Log2F.C. (IP/Input) for MYC mRNA. B. Genomic distribution of fRIP-seq peaks differentially enriched either in nucleoplasmic (NP) or chromatin (ca) fraction or distributed equally between chromatin and nucleoplasmic fractions. C. fRIP-PCR using TFF1 eRNA oligos. D. Summary plot showing the log2F.C.(fRIP IP/Input) across categories of ERα interacting RNA in Veh and E2 treatment. E. Heat map depicting ERα intensity on intergenic regions intersecting within 10kb of caRNA or NPRNA or ca=NP RNA enriched fRIP-seq peaks. F. Plot depicting the distance between the nearest ERα peak and fRIP-seq peak enriched either in caRNA, NPRNA or ca=NP RNA. Statistical significance determined by Mann Whitney U test. G. Illustration explaining the RNA mediated recruitment of ERα.

**Supplementary Figure S2.**
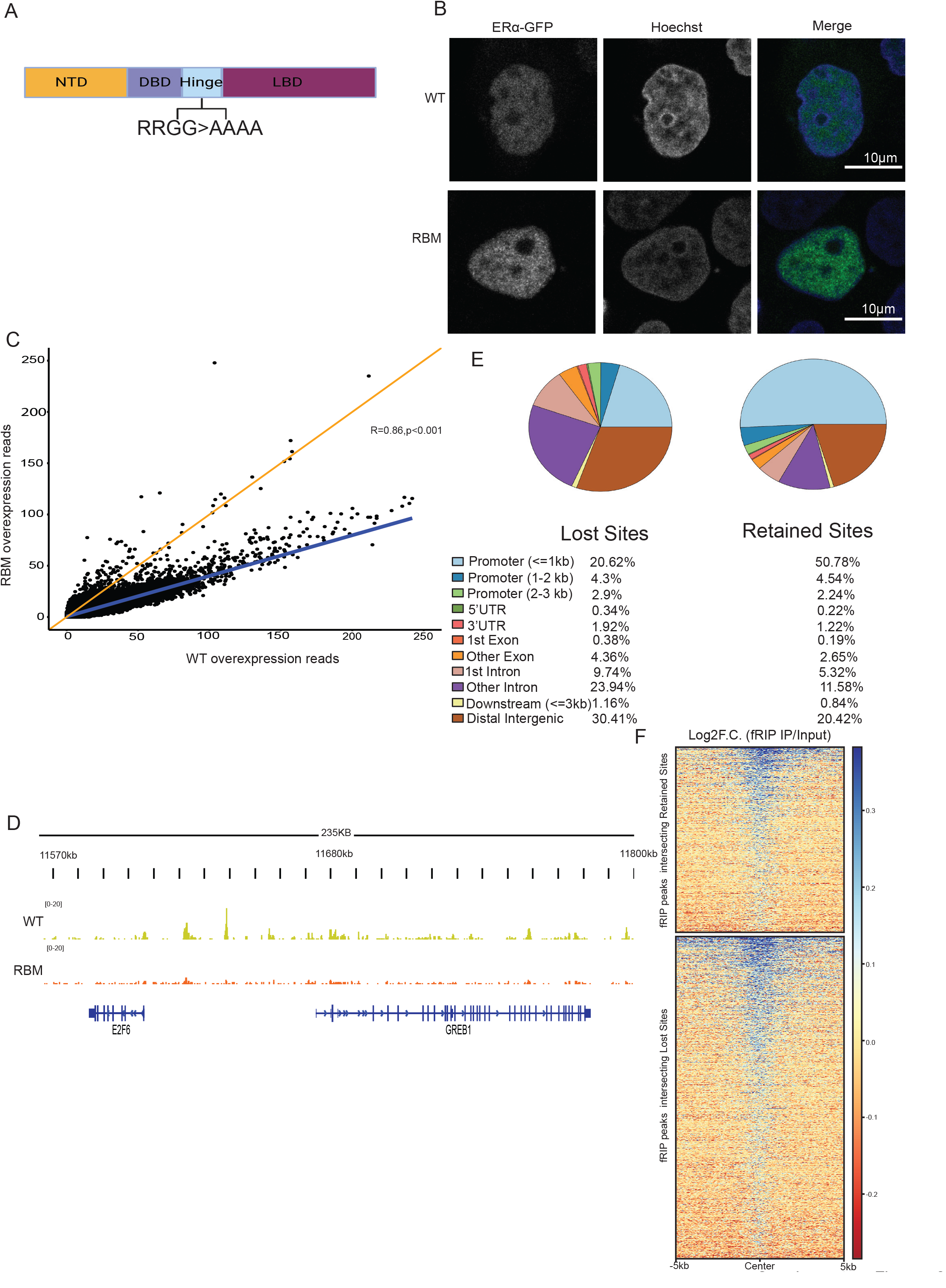
(Related to Figure 2) A. Schematic depicting the domains of ERα protein, the RNA binding RRGG sequence and its mutation to AAAA. B. Confocal images of ERα: GFP WT and RBM over-expressed in MCF-7 with E2 treatment for 1 hour. C. Normalized read counts showing the distribution of ERα:FLAG WT and RBM ChIP-seq reads. D. Genome browser snapshot showing ChIP-seq signal of ERα WT and RBM on GREB1 locus in MCF-7. E. Genomic distribution of lost and retained peaks upon over-expression of RBM-ERα over WT- ERα. F. Log2F.C.IP/Input for the fRIP-seq peaks intersecting with the lost and retained ERα upon RBM-ERα overexpression over WT-ERα.

**Supplementary Figure S3.**
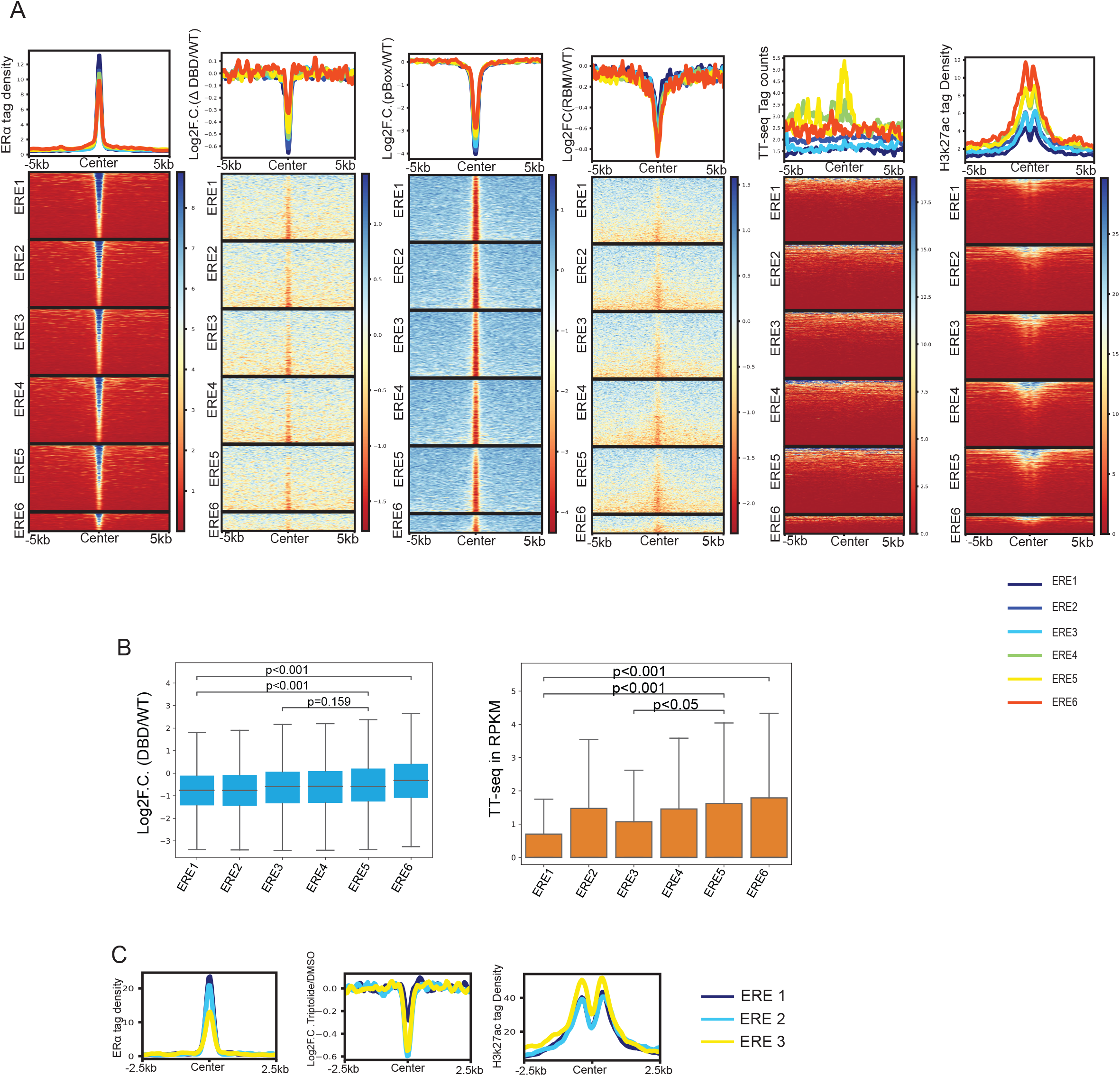
(Related to Figure 2) A. Heatmap depicting the ERα tag density, Log2F.C of (ΔDBD/WT), (pbox/WT), (RBM/WT), TT- seq (RPKM) and H3k27ac at ERE bins of varying strength (strongest to weakest) B. Boxplot depicting the Log2F.C of (ΔDBD/WT) and TT-seq at decreasing order of ERE strength. Statistical significance determined by Mann-Whitney U-test. C. Heat map depicting ERα tag density, Log2F.C (Triptolide ERα/DMSO ERα) tag density and H3K27ac tag density at varying ERE strength.

**Supplementary Figure S4.**
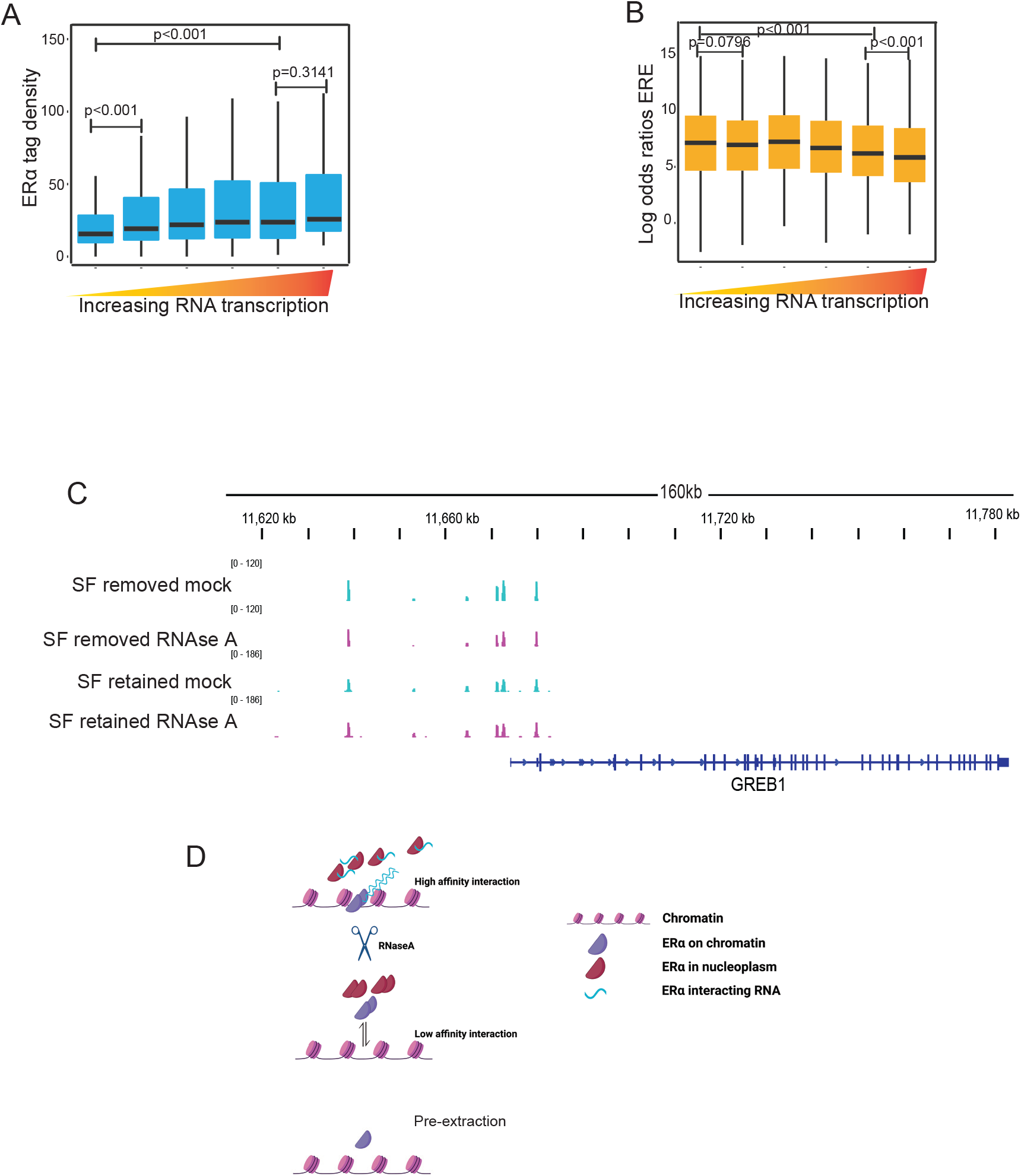
(Related to Figure 3) A. Boxplot showing ERα enrichment on all sites binned based on the levels of RNA transcription in increasing order. B. Boxplot showing the Log2 odds ratio for the ERE motif on all sites binned based on the levels of RNA transcription in increasing order. C. Genome Browser screenshot of GREB1 locus showing ERα ChIP-seq upon RNase A treatment with removal and retention of soluble proteins. D. Illustration depicting the RNA is required for ERα on chromatin.

**Supplementary Figure S5.**
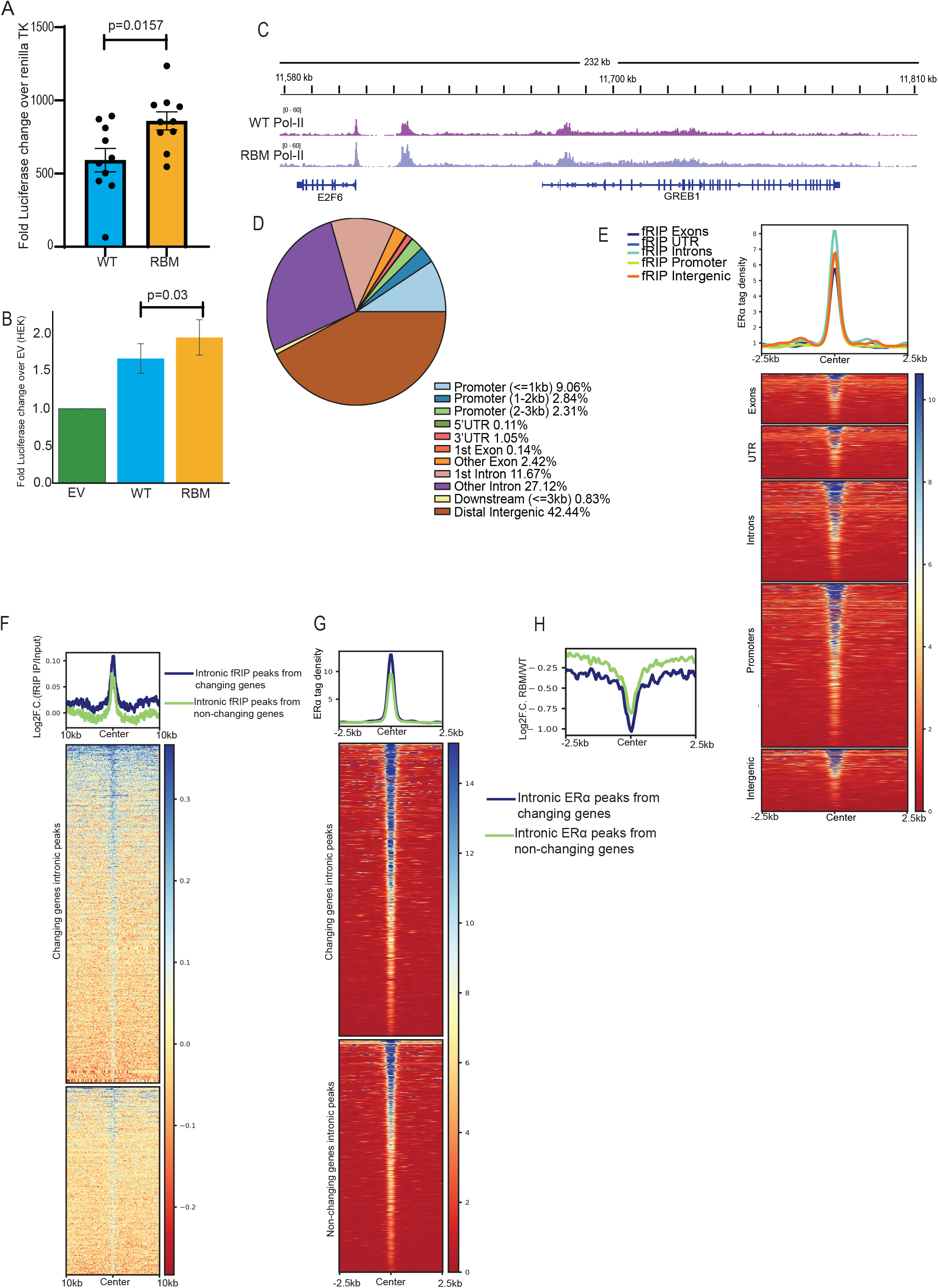
(Related to Figure 5) A. 3X ERE driven firefly luciferase activity normalized to renillaTK luciferase upon 24 hours of E2 treatment. Statistical significance determined by Mann-Whitney U-test. B. 3X ERE driven firefly luciferase activity upon WT-ERα or RBM-ERα overexpression and 24 hours of E2 treatment in HEK-293T. Statistical significance determined by Mann-Whitney U-test. C. Genome browser screenshot showing the occupancy of total PolII on GREB1 locus upon expression of WT-ERα or RBM-ERα. D. Pie chart depicting genomic distribution of ERα ChIP-seq peak. E. Heatmap depicting the ERα intensity across ERα bound regions within 10kb of various categories of fRIP-seq peaks. F. Heatmap depicting the fRIP-seq signal from intronic peaks of genes that are changing and non-changing. G. Heatmap depicting the ERα tag density on intronic sites from changing and non-changing gene categories. H. Profile plot depicting the log2FC (RBM-ERα/WT-ERα) for the intronic peaks from changing and non-changing gene categories.

**Table S1:**
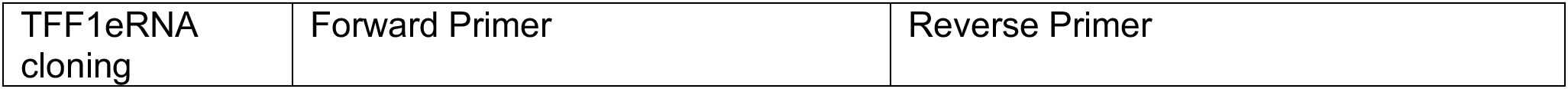
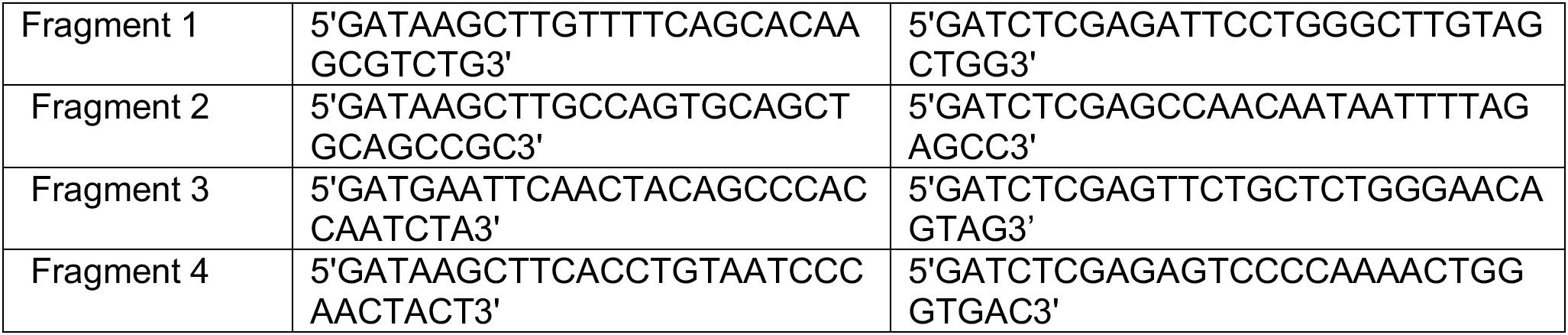
Primers used for cloning.

**TableS2:**
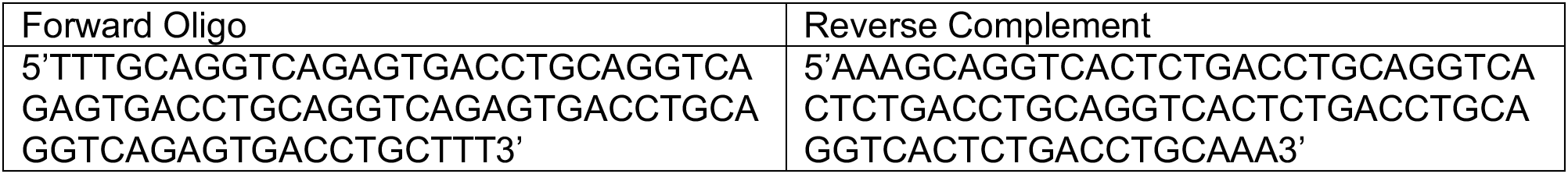
Sequence of 3X ERE Oligos.

**TableS3:**
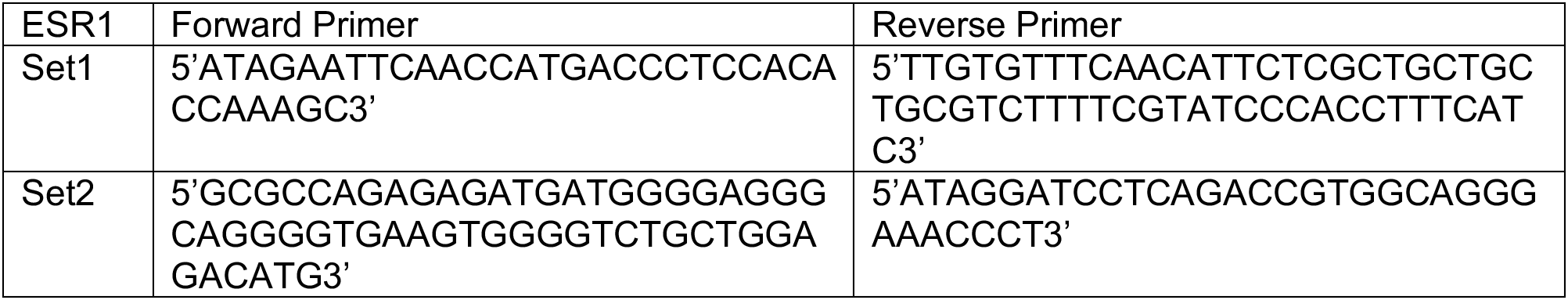
Primers used for cloning of RRGG>AAAA ESR1.

**TableS4:**
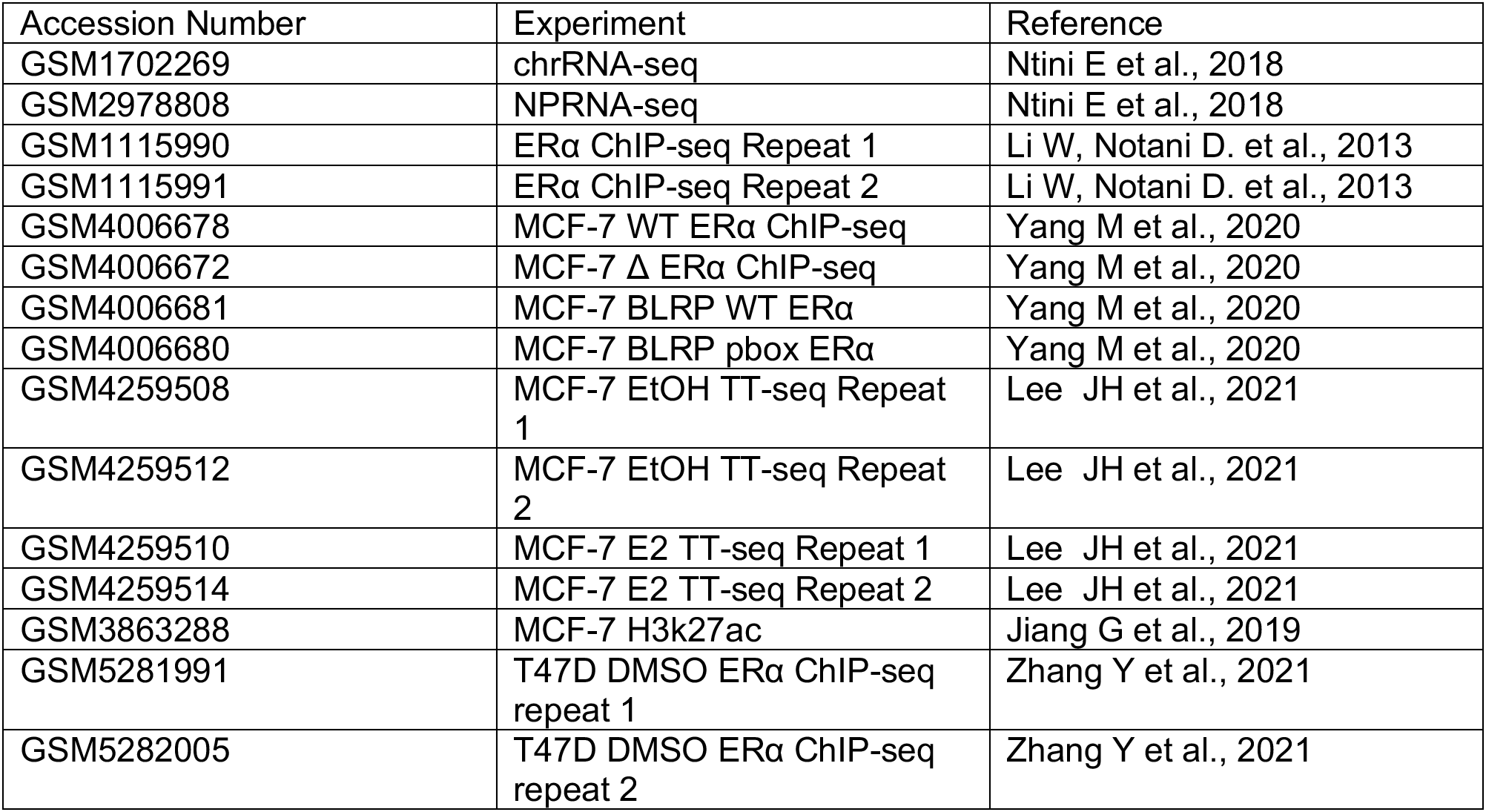
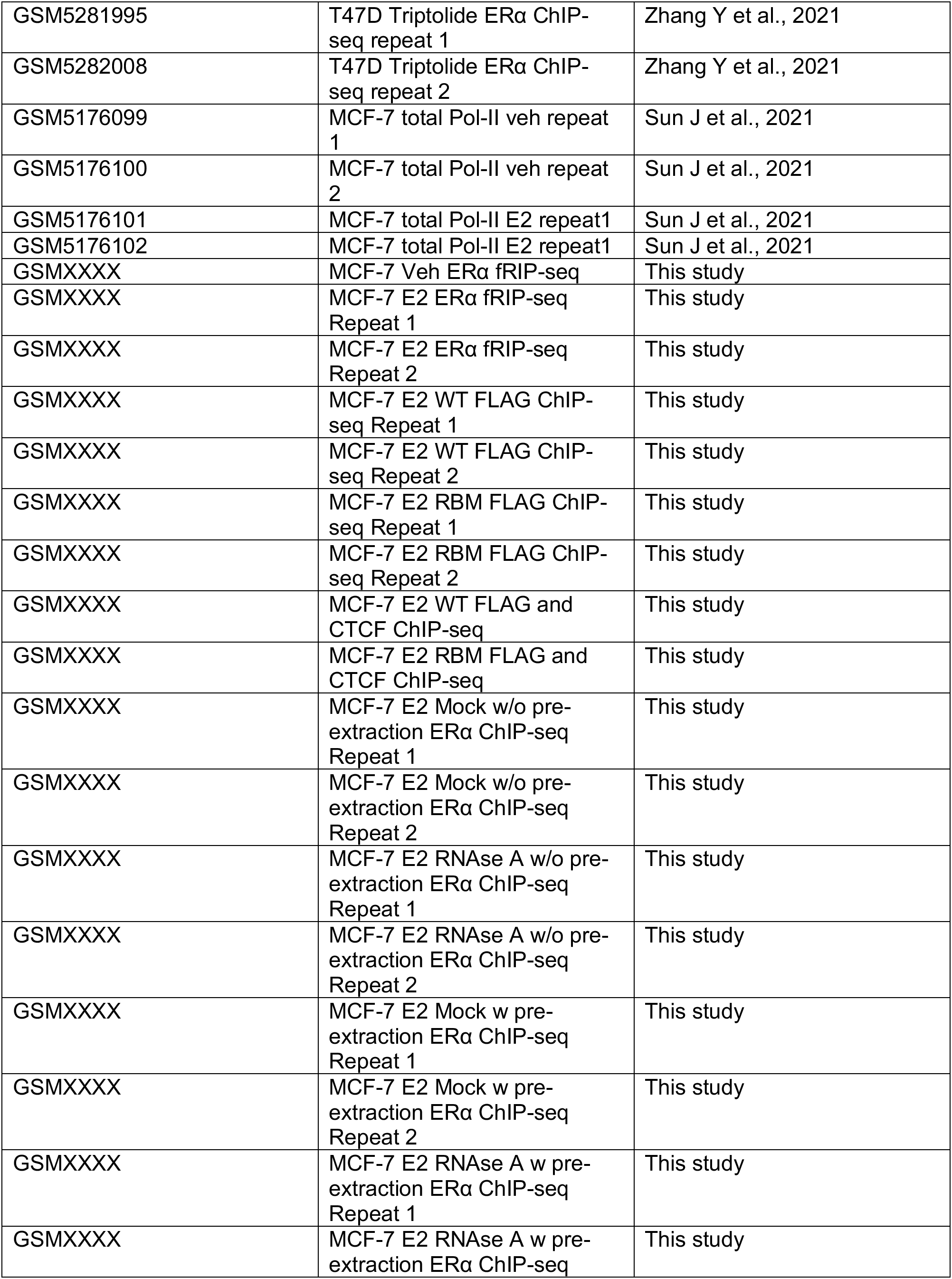
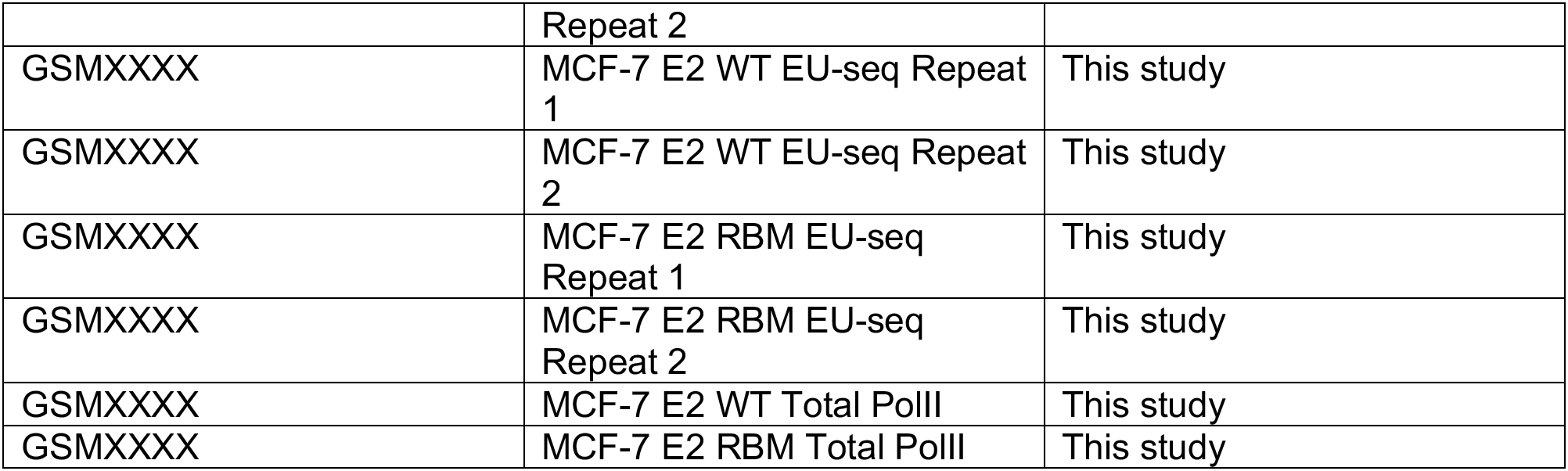
Accession number of NGS Datasets.

| Accession Number | Experiment | Reference |
| --- | --- | --- |
| GSM1702269 | chrRNA-seq | Ntini E et al., 2018 |
| GSM2978808 | NPRNA-seq | Ntini E et al., 2018 |
| GSM1115990 | ERα ChIP-seq Repeat 1 | Li W, Notani D. et al., 2013 |
| GSM1115991 | ERα ChIP-seq Repeat 2 | Li W, Notani D. et al., 2013 |
| GSM4006678 | MCF-7 WT ERα ChIP-seq | Yang M et al., 2020 |
| GSM4006672 | MCF-7 $\Delta$ ERα ChIP-seq | Yang M et al., 2020 |
| GSM4006681 | MCF-7 BLRP WT ERα | Yang M et al., 2020 |
| GSM4006680 | MCF-7 BLRP pbox ERα | Yang M et al., 2020 |
| GSM4259508 | MCF-7 EtOH TT-seq Repeat 1 | Lee JH et al., 2021 |
| GSM4259512 | MCF-7 EtOH TT-seq Repeat 2 | Lee JH et al., 2021 |
| GSM4259510 | MCF-7 E2 TT-seq Repeat 1 | Lee JH et al., 2021 |
| GSM4259514 | MCF-7 E2 TT-seq Repeat 2 | Lee JH et al., 2021 |
| GSM3863288 | MCF-7 H3k27ac | Jiang G et al., 2019 |
| GSM5281991 | T47D DMSO ERα ChIP-seq repeat 1 | Zhang Y et al., 2021 |
| GSM5282005 | T47D DMSO ERα ChIP-seq repeat 2 | Zhang Y et al., 2021 |
| GSM5281995 | T47D Triptolide ERα ChIP-seq repeat 1 | Zhang Y et al., 2021 |
| GSM5282008 | T47D Triptolide ERα ChIP-seq repeat 2 | Zhang Y et al., 2021 |
| GSM5176099 | MCF-7 total Pol-II veh repeat 1 | Sun J et al., 2021 |
| GSM5176100 | MCF-7 total Pol-II veh repeat 2 | Sun J et al., 2021 |
| GSM5176101 | MCF-7 total Pol-II E2 repeat1 | Sun J et al., 2021 |
| GSM5176102 | MCF-7 total Pol-II E2 repeat1 | Sun J et al., 2021 |
| GSMXXXX | MCF-7 Veh ERα fRIP-seq | This study |
| GSMXXXX | MCF-7 E2 ERα fRIP-seq Repeat 1 | This study |
| GSMXXXX | MCF-7 E2 ERα fRIP-seq Repeat 2 | This study |
| GSMXXXX | MCF-7 E2 WT FLAG ChIP-seq Repeat 1 | This study |
| GSMXXXX | MCF-7 E2 WT FLAG ChIP-seq Repeat 2 | This study |
| GSMXXXX | MCF-7 E2 RBM FLAG ChIP-seq Repeat 1 | This study |
| GSMXXXX | MCF-7 E2 RBM FLAG ChIP-seq Repeat 2 | This study |
| GSMXXXX | MCF-7 E2 WT FLAG and CTCF ChIP-seq | This study |
| GSMXXXX | MCF-7 E2 RBM FLAG and CTCF ChIP-seq | This study |
| GSMXXXX | MCF-7 E2 Mock w/o pre-extraction ERα ChIP-seq Repeat 1 | This study |
| GSMXXXX | MCF-7 E2 Mock w/o pre-extraction ERα ChIP-seq Repeat 2 | This study |
| GSMXXXX | MCF-7 E2 RNase A w/o pre-extraction ERα ChIP-seq Repeat 1 | This study |
| GSMXXXX | MCF-7 E2 RNase A w/o pre-extraction ERα ChIP-seq Repeat 2 | This study |
| GSMXXXX | MCF-7 E2 Mock w pre-extraction ERα ChIP-seq Repeat 1 | This study |
| GSMXXXX | MCF-7 E2 Mock w pre-extraction ERα ChIP-seq Repeat 2 | This study |
| GSMXXXX | MCF-7 E2 RNase A w pre-extraction ERα ChIP-seq Repeat 1 | This study |
| GSMXXXX | MCF-7 E2 RNase A w pre-extraction ERα ChIP-seq | This study |
|  | Repeat 2 |  |
| GSMXXXX | MCF-7 E2 WT EU-seq Repeat 1 | This study |
| GSMXXXX | MCF-7 E2 WT EU-seq Repeat 2 | This study |
| GSMXXXX | MCF-7 E2 RBM EU-seq Repeat 1 | This study |
| GSMXXXX | MCF-7 E2 RBM EU-seq Repeat 2 | This study |
| GSMXXXX | MCF-7 E2 WT Total PolII | This study |
| GSMXXXX | MCF-7 E2 RBM Total PolII | This study |

